# PPP2R1A Regulates Migration Persistence through the WAVE Shell Complex

**DOI:** 10.1101/2022.06.02.494622

**Authors:** Yanan Wang, Giovanni Chiappetta, Raphaël Guérois, Stéphane Romero, Matthias Krause, Claire Dessalles, Avin Babataheri, Abdul I. Barakat, Joelle Vinh, Anna Polesskaya, Alexis M. Gautreau

**Author notes:** These authors contributed equally to the work (Giovanni Chiappetta, Raphaël Guérois, Stéphane Romero).

## Abstract

The RAC1-WAVE-Arp2/3 signaling pathway generates branched actin networks that power lamellipodium protrusion of migrating cells. Feedback is thought to control protrusion lifetime and migration persistence, but its molecular circuitry remains elusive. Using proteomics, we identified PPP2R1A among proteins differentially associated with the WAVE complex subunit ABI1 when RAC1 was activated and downstream generation of branched actin was blocked. PPP2R1A was found to associate at the lamellipodial edge with a novel form of WAVE complex, the WAVE Shell Complex (WSC), that contains NHSL1 instead of the Arp2/3 activating subunit WAVE as in the canonical WAVE Regulatory Complex (WRC). PPP2R1A was required for persistence in random and directed migration assays and for RAC1-dependent actin polymerization in cell extracts. PPP2R1A requirement was abolished by NHSL1 depletion. PPP2R1A mutations found in tumors impaired WSC binding and migration regulation, suggesting that this novel function of PPP2R1A is critical for its tumor suppressor activity.

---

Cell migration is a critical process for animal cells, especially in the embryo, but also in the adult. For example, immune cells constantly patrol the organism to fight infections. Nucleation of branched actin by the Arp2/3 complex fuels membrane protrusions called lamellipodia. This type of cell migration is characterized by its persistence that can be seen even in random migration and measured by the time during which direction is maintained once it is established ^1^. Arp2/3-mediated persistence also favors directed migration towards higher concentrations of extracellular matrix (ECM) proteins in a process called haptotaxis ^2^.

Three WAVE proteins activate Arp2/3 at the cell cortex and in membrane protrusions ^3^. WAVE proteins are subunits of the WAVE Regulatory Complex (WRC) that maintains WAVE inactive ^4,5^. Upon binding to GTP-bound RAC1, WRC activates Arp2/3 ^6,7^. The mechanism involves the exposure of the WCA domain of WAVE that binds and activates Arp2/3 ^8,9^. The RAC1-WAVE-Arp2/3 pathway is critical for development and normal adult life but is also involved in cancer ^10–12^. Genes encoding subunits of the WAVE and Arp2/3 complexes are overexpressed in a variety of cancers, and this overexpression is associated with high tumor grade and poor patient prognosis ^13^.

Cell migration is finely regulated at all molecular levels. Each positive component required to generate cortical branched actin, RAC1, WRC and Arp2/3, appears to be counteracted by inhibitory proteins, CYRI ^14,15^, NHSL1 ^16^ and ARPIN ^17,18^, respectively. NHSL1 belongs to the family of Nance-Horan Syndrome (NHS) proteins, which contain an N-terminal WAVE Homology Domain (WHD), as in WAVE proteins ^19^. The WHD is the main structural domain that embeds WAVE proteins in the WRC ^6,9^, raising the possibility that WRC might contain NHS family proteins instead of WAVE proteins at some point in their life cycle. This intriguing possibility, however, has never been reported until now. Instead NHSL1 has been shown to interact with WRC through the C-terminal SH3 domain of the WRC subunit ABI1 ^16^.

Regulatory circuits of cell migration involve feedback and feedforward loops ^20,21^. ARPIN was shown to inhibit Arp2/3 only when RAC1 signaling was on, rendering lamellipodia unstable once they were formed instead of preventing their formation ^17^. Positive feedback that sustains membrane protrusion at the front is thought to be responsible for the persistence of cell migration. Indeed, signaling pathways are not linear, and actin polymerization activate WRC further in space and time in propagating waves ^22,23^ and promotes WRC turn-over ^24^. Biochemical signaling in feedback is constrained by cell mechanics and in particular membrane tension, which appears as a central component to sustain lamellipodial protrusion at the leading edge ^25,26^. Feedback signaling is so challenging to dissect that mathematical modeling, computational simulations and machine learning are often required to interpret complex observations that would not fit into simple and linear models ^27,28^.

Here we used criteria of feedback signaling to uncover a novel molecular machine that regulates migration persistence. Using differential proteomics, we identified the PPP2R1A protein as a novel migration regulator and found that its mode of action involves an interaction with a new multiprotein assembly containing NHSL1 in place of a WAVE subunit, the WAVE Shell Complex (WSC).

## RESULTS

### Identification of PPP2R1A as a Novel Regulator of Migration Persistence

To identify novel partners of WRC, we used proteomics after tandem affinity purification (TAP) of the ABI1 subunit. We reasoned that partners differentially associated with ABI1 when RAC1 is activated and when Arp2/3 is inhibited would be good candidates to regulate migration persistence and feedback signaling. As a model system, we used the human mammary epithelial cell line MCF10A, which is immortalized but not transformed. We thus isolated stable cell lines expressing FLAG-GFP ABI1 from both parental and a genome-edited derivative, where the Q61L mutation affects one allele of *RAC1* and renders the RAC1 GTPase deficient and hence constitutively active. MCF10A cells expressing RAC1 Q61L showed more persistent migration than parental cells ^29^. We treated or not both cell lines with the Arp2/3 inhibitory compound CK-666 to modulate the binding of potential WRC partners involved in signaling feedback from branched actin.

After the two successive FLAG and GFP immunoprecipitations (Fig.1a), we identified by mass spectrometry 89 proteins specifically associated with ABI1, but not with FLAG-GFP control (Table S1). We performed label-free quantification of triplicates and calculated the relative abundance of associated proteins in all four conditions after normalization by the amount of the ABI1 bait protein retrieved in each condition. The WRC subunits did not vary considerably in the different conditions, 20 % at most (Fig.1b). Among the hits, we recognized several described functional partners of the WRC such as SRGAP3, also known as WRP ^30^, the Nance-Horan Syndrome family proteins NHS and NHSL1 ^16,19^, IRSp53 ^31^ and lamellipodin, which was 3-fold more associated with WRC when RAC1 was activated, as previously reported ^32^.

**Figure 1.**
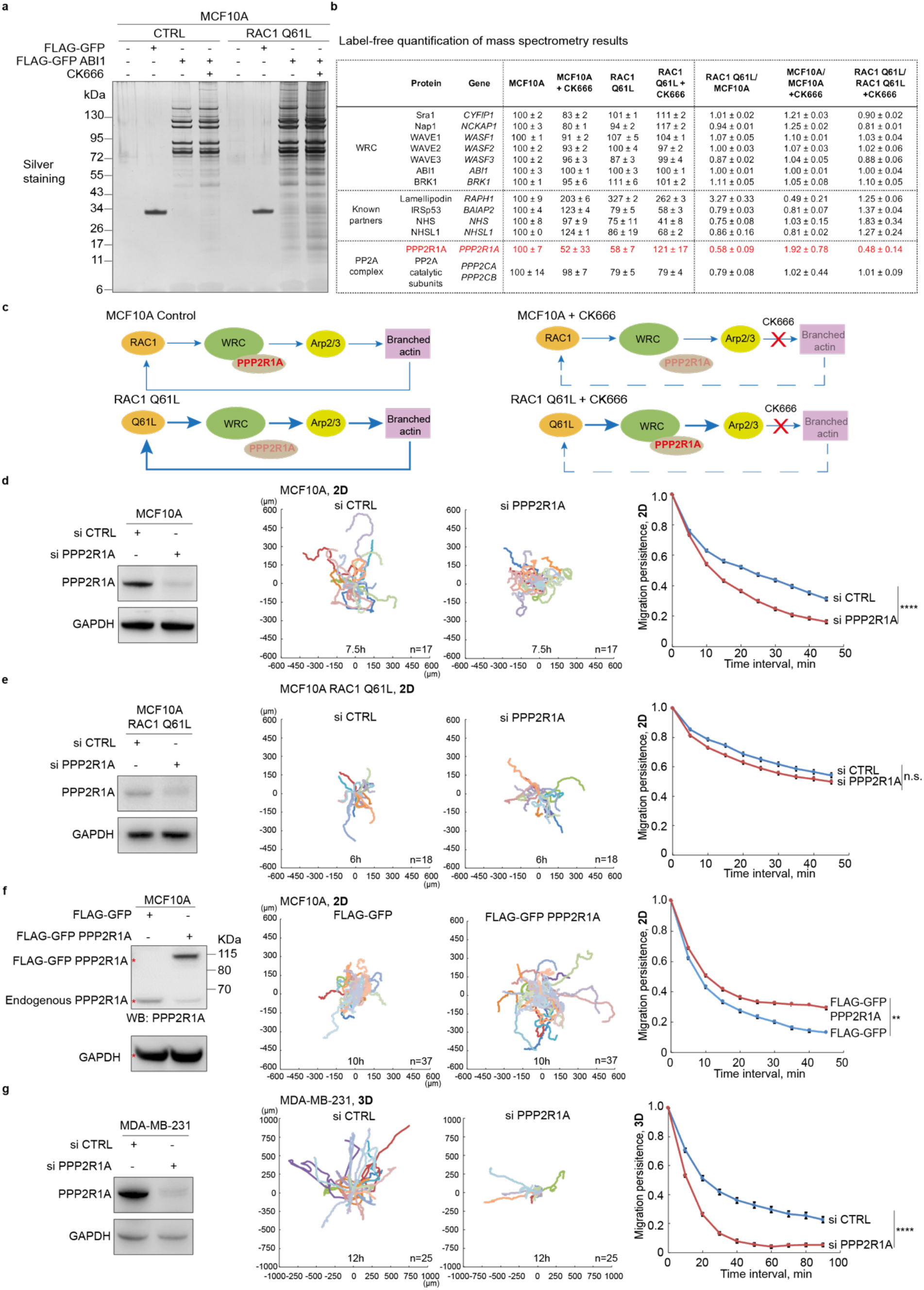
Identification of PPP2R1A as an ABI1 partner that regulates migration persistence. **(a)** MCF10A parental cells or genome-edited MCF10A cells where a *RAC1* allele encodes the active RAC1 Q61L mutant form were stably transfected with plasmids expressing FLAG-GFP or FLAG-GFP ABI1. Cells were treated or not with CK666 in order to block the polymerization of branched actin and thus the suspected feedback loop that mediates migration persistence. FLAG-GFP proteins were then purified by Tandem Affinity Purification and associated proteins were revealed by silver staining of SDS-PAGE gels. **(b)** PPP2R1A associates with ABI1 differently upon RAC1-dependent Arp2/3 activation. Label-free quantification of selected proteins identified by mass spectrometry to be associated with ABI1. 3 technical repeats, mean ± sem. **(c)** Schematic representations of the association of PPP2R1A with the WRC in different conditions. **(d)** MCF10A cells were transfected with pools of control (CTRL) or PPP2R1A siRNAs and analyzed by Western blots with PPP2R1A and GAPDH antibodies. Cell trajectories and migration persistence extracted from 2D migration of single MCF10A cells transfected with indicated siRNAs. **(e)** Same experiment as in (a) with MCF10A RAC1 Q61L cells. **(f)** MCF10A cells were stably transfected with plasmids expressing FLAG-GFP or FLAG-GFP PPP2R1A. Migration was analyzed as in (a). **(g)** Same experiment as in (a) with MDA-MB-231 cells analyzed in collagen type I gels. Most cell trajectories upon PPP2R1A depletion are so short that they cannot be well distinguished, because they overlap in the center of the graph. 3 biological repeats of each experiment yielded similar results, only one is displayed. **P<0.01; ****P<0.0001; n.s. not significant.

Our attention was drawn to PPP2R1A, a scaffold subunit of the PP2A phosphatase complex, which showed variations in the two-fold range when RAC1 was activated and when CK-666 was applied to cells. The amount of PPP2R1A associated with ABI1 decreased when RAC1 was constitutively activated (Fig.1c). CK-666 induced opposite PPP2R1A variations in parental and RAC1 Q61L cells, probably because the constitutively active RAC1 Q61L does not depend on the feedback of branched actin for its activity. Catalytic subunits of the same PP2A phosphatase complex, encoded by the two paralogous genes, *PPP2CA* and *PPP2CB*, were also detected in the list of ABI1 partners, but surprisingly these subunits did not show the same variations as PPP2R1A when CK-666 was applied to cells.

We depleted PPP2R1A from MCF10A cells using siRNA pools and tested cells in a 2D random migration assay. Lamellipodia appeared less developed in PPP2R1A-depleted cells, which, as a result, were less spread than control cells (Movie S1). PPP2R1A-depleted cells did not maintain the direction of migration over time as well as controls (Fig.1d). In contrast, this decrease in migration persistence upon PPP2R1A depletion was not significant when the same assay was performed using MCF10A cells expressing RAC1 Q61L (Fig.1e, Movie S2). This was surprising given that migration persistence is increased in MCF10A RAC1 Q61L ^29^, but was in agreement with our observation that PPP2R1A interacts less with ABI1 when RAC1 is constitutively active. We then isolated a stable MCF10A cell line that expresses FLAG-GFP PPP2R1A. The overexpression of tagged PPP2R1A was moderate, estimated by densitometry of PPP2R1A Western blots to be less than two-fold (1.96 ± 0.32, mean ± sem of 3 triplicates, P < 0.05, t-test), with a concomitant decrease in endogenous PPP2R1A (Fig.1f). PPP2R1A expression was, however, sufficient to increase migration persistence in the 2D random migration assay (Movie S3). The loss- and gain-of-function experiments in MCF10A cells thus indicate that migration persistence depends on PPP2R1A levels.

To validate our findings in another cell system, we used the invasive breast cancer cell line MDA-MB-231 that we embedded in 3D collagen gels. PPP2R1A depletion dramatically decreased migration persistence in this setting as well (Fig.1g, Movie S4). Overexpression of PPP2R1A, however, did not affect migration persistence in either MDA-MB-231 cells, or in MCF10A cells expressing RAC1 Q61L (Fig.S1). Migration persistence of cells expressing constitutively active RAC1 Q61L thus appears independent of PPP2R1A levels, unlike parental MCF10A. Other parameters of cell migration necessary for a complete description of cell trajectories, such as cell speed and Mean Square Displacement (MSD), were also altered in some of these genetic perturbations, but inconsistently (Fig.S1), as previously reported ^17,29^, confirming that migration persistence, but not these other parameters, is the primary target of the RAC1-WAVE-Arp2/3 pathway ^20^.

### PPP2R1A Interacts With the NHSL1-containing WAVE Shell Complex

To decipher how PPP2R1A regulated migration of MCF10A and MDA-MB-231 cells, we characterized its partners by proteomics. We purified FLAG-GFP-PPP2R1A by TAP (Fig.2a) and identified its partners by mass spectrometry (Table S2). As expected, we detected in MCF10A and MDA-MB-231 cells the catalytic phosphatase subunit and numerous regulatory subunits of PP2A complexes (PPP2R1B, PPP2R2D, PPP2R3B, PPP2R5A, PPP2R5B, PPP2R5C, PPP2R5D, PPP2R5E) as well as its larger STRIPAK derivatives (STRN, STRIP2, STRN4, MOB4). PPP2R3C was detected associated with PPP2R1A in MCF10A, but not MDA-MB-231 cells. We expected to find the whole WRC in line with the TAP purification of ABI1. However, we detected only 4 of the 5 subunits of the WRC (Fig.2b), namely CYFIP1, NCKAP1, ABI1 and BRK1. BRK1 was only detected in MCF10A and not in MDA-MB-231 cells, but this is likely because BRK1 generates only a few tryptic peptides from its short sequence of 75 amino-acids. Surprisingly, the 3 WAVE proteins were absent from the TAP purification from both cell lines. We searched for NHS family proteins, which were reported to contain an N-terminal WAVE homology domain (WHD) similar to that responsible for WAVE incorporation into WRC ^6,19^, and found NHSL1 in the TAP purification from both cell lines. We validated the presence of NHSL1 and all WRC subunits except WAVE by Western blots (Fig.2c). We never detected any WAVE protein in TAP purification of PPP2R1A. When NHSL1 was depleted from cells using siRNAs, PPP2R1A immunoprecipitation did not retrieve WRC subunits, but still retrieved the PP2A catalytic subunit (Fig.2d). This result suggests that NHSL1 is the subunit of this alternative complex that is recognized by PPP2R1A.

**Figure 2.**
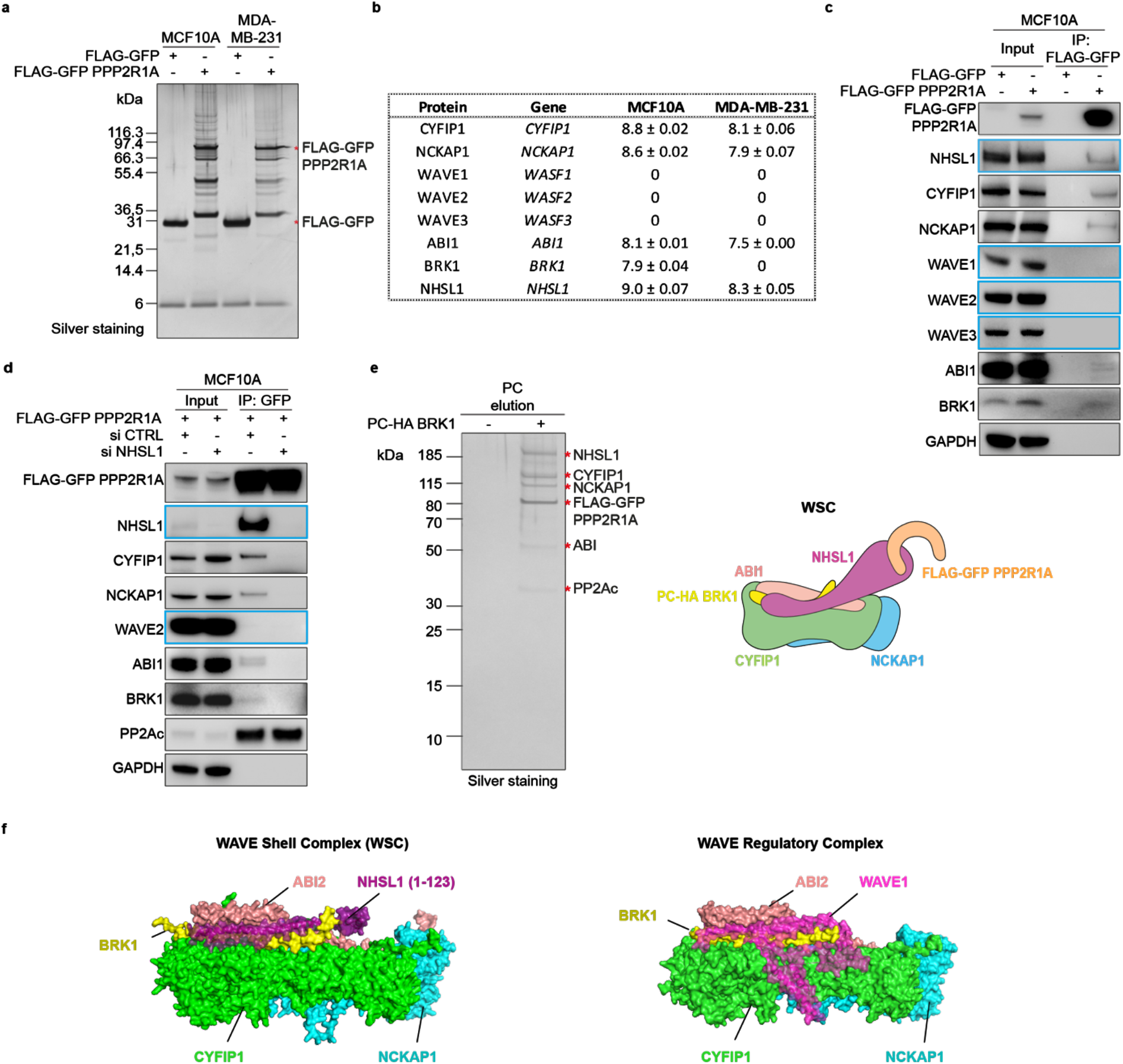
PPP2R1A interacts with the WAVE Shell Complex (WSC) that contains NHSL1 instead of WAVE proteins. **(a)** MCF10A or MDA-MB-231 cells stably expressing FLAG-GFP or FLAG-GFP PPP2R1A were subjected to FLAG-GFP Tandem Affinity Purification and purified proteins were resolved by SDS-PAGE and silver stained. The same samples were used for mass spectrometry and Western blots. **(b)** Label-free quantification of proteins identified by mass spectrometry. In both cell lines, NHSL1 is present at the expense of WAVE subunits. 3 technical repeats, mean ± sem. **(c)** Western blots with the indicated antibodies. 3 biological repeats with similar results. **(d)** PPP2R1A associates with the WSC through NHSL1. MCF10A cells expressing FLAG-GFP PPP2R1A were transfected with pools of siRNAs targeting NHSL1 or non-targeting siRNAs (CTRL). GFP immunoprecipitates were analyzed by Western blots. 3 biological repeats with similar results. **(e)** An MCF10A cell line stably transfected with two plasmids expressing FLAG-GFP PPP2R1A and PC-HA BRK1 was subjected to FLAG-PC Tandem Affinity Purification to purify the WSC. WSC composition was analyzed by mass spectrometry. Silver stained SDS PAGE of purified WSC. The identity of WSC subunits and the lack of WAVE2 was confirmed using Western blots. 3 biological repeats with similar results. **(f)** Comparison of the structural model of the WSC obtained using Alphafold2 with the X-ray crystal structure WAVE Regulatory Complex (PDB:3P8C). The WSC was composed of NHSL1 (1-123) (purple), CYFIP1 (green), NCKAP1 (cyan), BRK1 (yellow) and ABI2 (1-160) (salmon).

To examine the composition of the alternative complex devoid of WAVE, we performed TAP to select for the presence of both PPP2R1A and the WRC subunit BRK1. We performed sequential FLAG - PC immunoprecipitations from the stable MCF10A cell line expressing both FLAG-GFP-PPP2R1A and PC-HA-BRK1. We indeed retrieved the WRC without WAVE but containing NHSL1 instead (Fig.2e, Table S3). This TAP purification of both PPP2R1A and BRK1 did not completely exclude PP2A subunits, since it also contained a subset of the subunits found in TAP of PPP2R1A alone, namely PPP2CA, PPP2R2A, PPP2R5D, PPP2R5E and STRN. This result suggests that PPP2R1A can bind to this alternative WRC while being part of PP2A complexes. To our knowledge, a physiological form of WRC without a WAVE subunit and containing an NHS family protein instead, has not previously been reported. NHSL1 was recently reported to interact with the complete canonical WRC through the ABI1 SH3 domain ^16^. PPP2R1A thus does not bind to all pools of NHSL1 molecules, but appears to select a specific conformation of NHSL1, where NHSL1 fully replaces WAVE within its complex. We call this alternative to WRC, the WAVE Shell Complex (WSC), to emphasize that this multiprotein complex, which normally embeds WAVE and regulates its activity, has lost its WAVE subunit.

The HHpred program that detects remote homology ^33^ indeed predicted with a high probability (99.9 %) that NHSL1 contains a WAVE homology domain (WHD) in its N-terminus (residues 30-90; Fig.S2). A high-confidence structural model generated by the AlphaFold2 program ^34–36^ showed how NHSL1 could interact with the CYFIP1, NCKAP1, BRK1 and ABI2 subunits (Fig.2f). In this model, NHSL1 forms a triple coiled-coil with BRK1 and ABI2 as the N-terminal helix of the WAVE1 subunit in the WRC ^6,37^. Unlike WAVE1 in the WRC, no other region of NHSL1, beyond the Pro97, is predicted to interact with WSC subunits. Three hydrophobic residues of NHSL1, upstream of its WHD, interact with CYFIP1 and contribute to the WSC assembly (Fig.S2).

To examine whether the PP2A complex can dephosphorylate WSC, or WRC, we performed TAP purification of ABI1 in the presence or absence of PPP2R1A (Fig.S3) and analyzed phosphosites by mass spectrometry. We detected phosphosites in ABI1, WAVE2, NHS, NHSL1 and IRSp53, among the well-characterized proteins of the pathway (Table S4). Label-free quantification of phosphosites is not as reliable as that of proteins, which are quantified by several peptides. Nevertheless, we quantified phosphosites when the WSC was connected to PP2A complexes via PPP2R1A or not. All but one phosphosite were either present in similar amounts in both conditions or decreased in the absence of PPP2R1A, which is inconsistent with the lack of association with a phosphatase in this condition. Only the phosphorylation of WAVE2 on Ser308 was found to be increased by 2.6-fold upon PPP2R1A depletion, as if this phosphosite was dephosphorylated by PP2A complexes. This phosphorylated residue, however, was not consistently detected in all technical repeats of the experiment. We attempted to validate this variation in phosphorylation using phosphospecific antibodies, but no increase in phosphorylation was detected in the absence of PPP2R1A (Fig.S3). The variation in the phosphorylation of WAVE2 Ser308 was in any case inconsistent with PP2A complexes associating with WSC, since WAVE2 is present in the WRC but not in the WSC. Thus, despite the presence of WSC-associated PP2A subunits, including the catalytic subunit, we found no evidence for WSC-or even WRC-directed phosphatase activity.

We then sought to examine PPP2R1A localization and used for this purpose B16-F1 cells that develop prominent lamellipodia. Cells were transfected with GFP-NHSL1 and mScarlet-PPP2R1A. PPP2R1A was colocalized with NHSL1 at the lamellipodium edge (Fig.3a, Movie S5). NHSL1 was strongly enriched at the lamellipodial edge, whereas PPP2R1A extended far beyond, in the entire width of the lamellipodium (Fig.3b). Arp2/3 is incorporated into branched actin networks and extends backwards through the actin retrograde flow. We thus examined the colocalization of mScarlet-PPP2R1A with the GFP-tagged Arp2/3 subunit ARPC1B. Both proteins colocalized in the width of lamellipodia (Fig.3c, Movie S6), but ARPC1B distribution declined steeply away from the edge, whereas PPP2R1A did not exhibit such a gradient (Fig.3d). These analyses suggest that PPP2R1A is a lamellipodial protein that is likely to interact with WSC at the cell edge.

**Figure 3.**
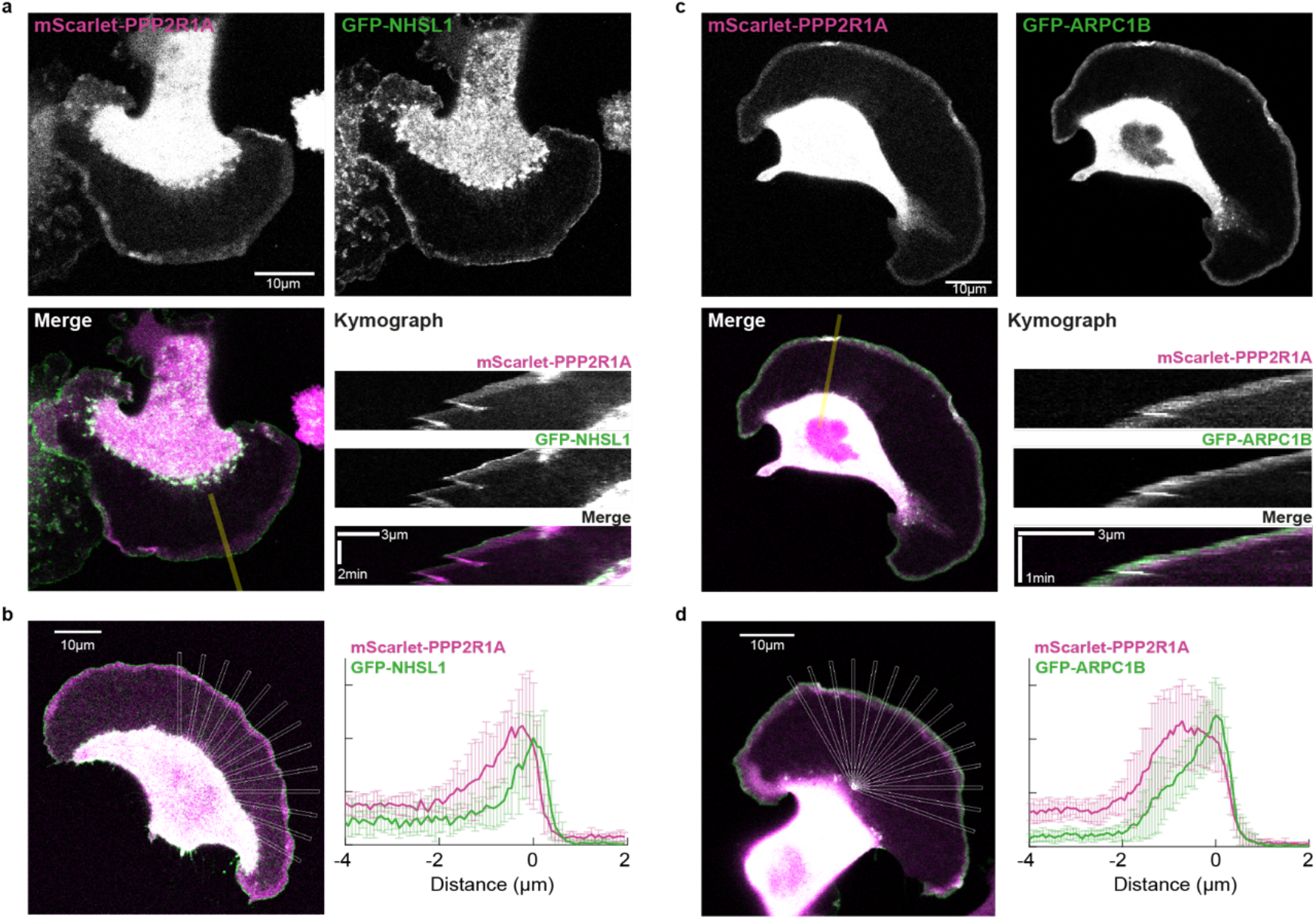
PPP2R1A colocalizes with branched actin in the width of the lamellipodium and colocalizes with NHSL1 at the lamellipodium edge. **(a)** B16-F1 cells were transiently transfected with mScarlet-PPP2R1A and GFP-NHSL1. A kymograph was drawn along the line shown on the merge. **(b)** Overlap of PPP2R1A and NHSL1 over multiple radial line scans registered to the cell edge. **(c)** B16-F1 cells were transiently transfected with mScarlet-PPP2R1A and the Arp2/3 subunit GFP-ARPC1B. A kymograph was drawn along the line shown on the merge. **(d)** Overlap of PPP2R1A and ARPC1B over multiple radial line scans registered to the cell edge.

### The Requirement of PPP2R1A in Cell Migration and Actin Polymerization Depends on the Presence of NHSL1

NHSL1 was previously described as a negative regulator of cell migration ^16^. It was thus surprising to find PPP2R1A and NHSL1 biochemically associated. We have knocked down either PPP2R1A, or NHSL1, as well as both proteins simultaneously in order to study epistasis. In line with previous findings, NHSL1 depletion strongly promoted migration persistence in random migration of single MCF10A cells. The decline of migration persistence upon PPP2R1A depletion was abolished by NHSL1 depletion (Fig.4a, Movie S7). Despite their opposite effects, neither of the two migration regulators appears to take over. The simultaneous depletion of both regulators does not significantly affect migration persistence compared to controls, as if the signaling circuit containing the PPP2R1A and NHSL1 was simply dispensable in this random migration assay. We then examined directional migration in a gradient of fibronectin to further challenge cells. This haptotactic assay was previously shown to depend on Arp2/3 activity ^2^. PPP2R1A-depleted cells were strongly impaired in their ability to migrate toward higher concentrations of fibronectin (Fig.4b, Movie S8). In contrast, depletion of the migration inhibitor NHSL1 rendered cells slightly, but significantly, more efficient at directed migration. In this assay, as in random migration, simultaneous depletion of PPP2R1A and NHSL1 abolished the effects of single depletions. In two different cell migration assays, PPP2R1A and NHSL1 thus depend on each other.

**Figure 4.**
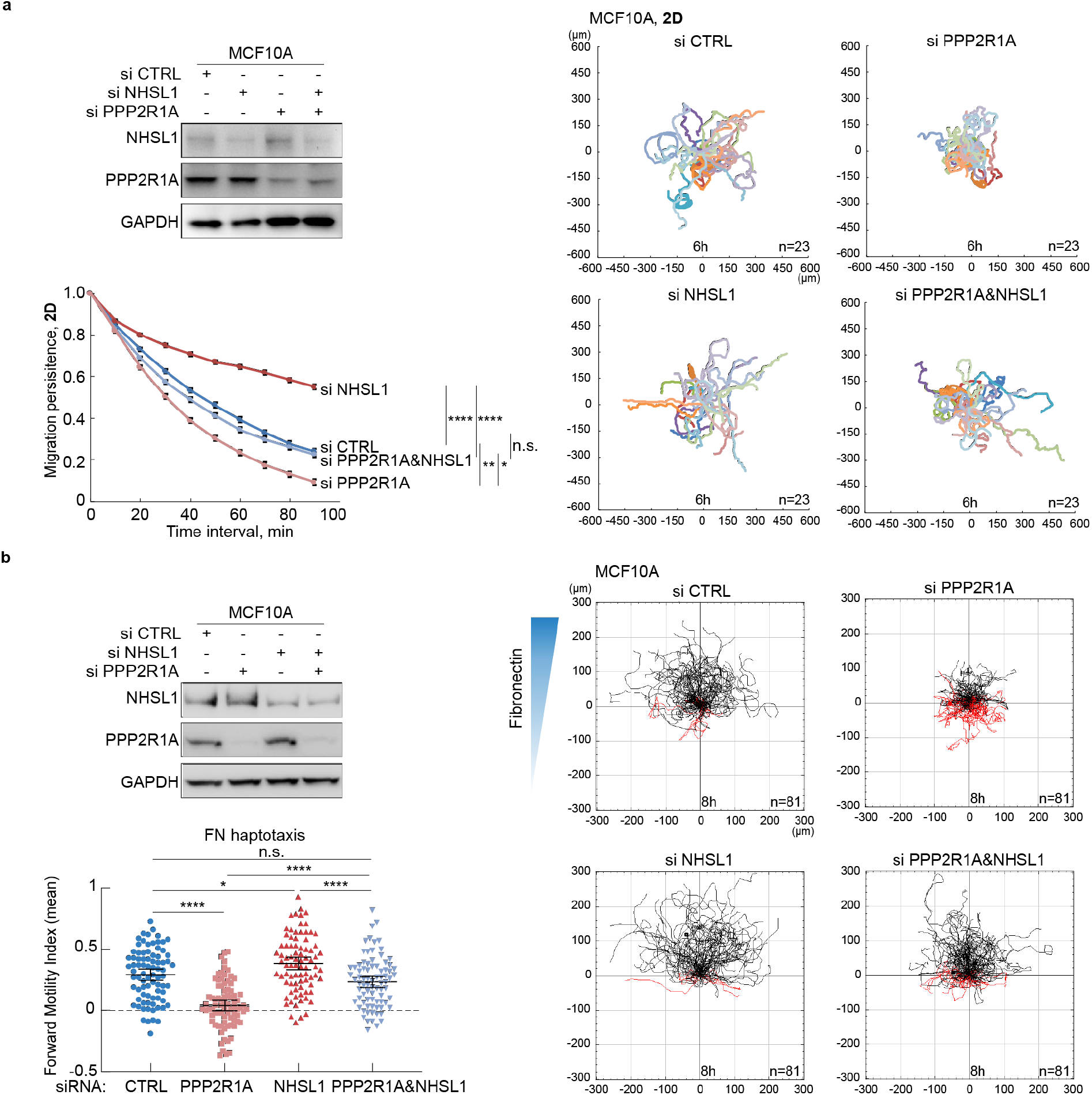
PPP2R1A depends on NHSL1 to regulate cell migration. MCF10A cells were transfected with pools of siRNAs targeting PPP2R1A, NHSL1 or both. Non-targeting siRNAs were transfected as controls (CTRL). Western blots were used to validate depletion. **(a)** 2D cell trajectories and migration persistence of single MCF10A cells transfected with indicated siRNAs, n=23. 3 biological repeats with similar results, only one is displayed. **(b)** Haptotaxis of MCF10A cells along a gradient of fibronectin. Cell trajectories of MCF10A cells transfected with indicated siRNAs. 3 biological repeats with similar results. All tracked cells from the 3 experiments were pooled to reach n=81 cells. Mean ± 95% confidence intervals of Forward Motility Index are plotted. *P<0.05; **P<0.01; ****P<0.0001

To establish that the effects of the PPP2R1A-NHSL1 signaling circuit were due to actin polymerization, we set up an in vitro reconstitution assay of actin polymerization in extracts prepared from MCF10A cells. Actin polymerization at the surface of glutathione beads was monitored by the incorporation of fluorescent actin. GTP-bound RAC1 Q61L was able to trigger the polymerization of actin structures from the surface of the beads (Fig.5a). These structures were discrete fibers, not the dense meshwork observed around beads displaying CDC42 Q61L (Fig.S4). These actin structures, however, were composed of branched actin, since their formation was impaired by the CK-666 compound, as measured by their total fluorescence and length. These actin structures depended on the WRC, as shown using extracts prepared from NCKAP1-depleted cells, and did not depend on the CDC42 effector N-WASP, since they were not inhibited by the N-WASP inhibitory compound wiskostatin ^38^ (Fig.S4). Extracts prepared from PPP2R1A-depleted cells did not support actin polymerization from RAC1 Q61L displaying beads (Fig.5b). Yet the extracts prepared from NHSL1-depleted cells and those prepared from cells simultaneously depleted of PPP2R1A and NHSL1 were as competent as wild type extracts in supporting actin polymerization. Both PPP2R1A and NHSL1 were detected associated with GTP-bound RAC1 Q61L (Fig.S4), in line with the fact that both CYFIP1 and NHSL1 harbor binding sites for active RAC1 ^7,16^. Together, these results show that PPP2R1A requires NHSL1 to regulate cell migration and actin polymerization.

**Figure 5.**
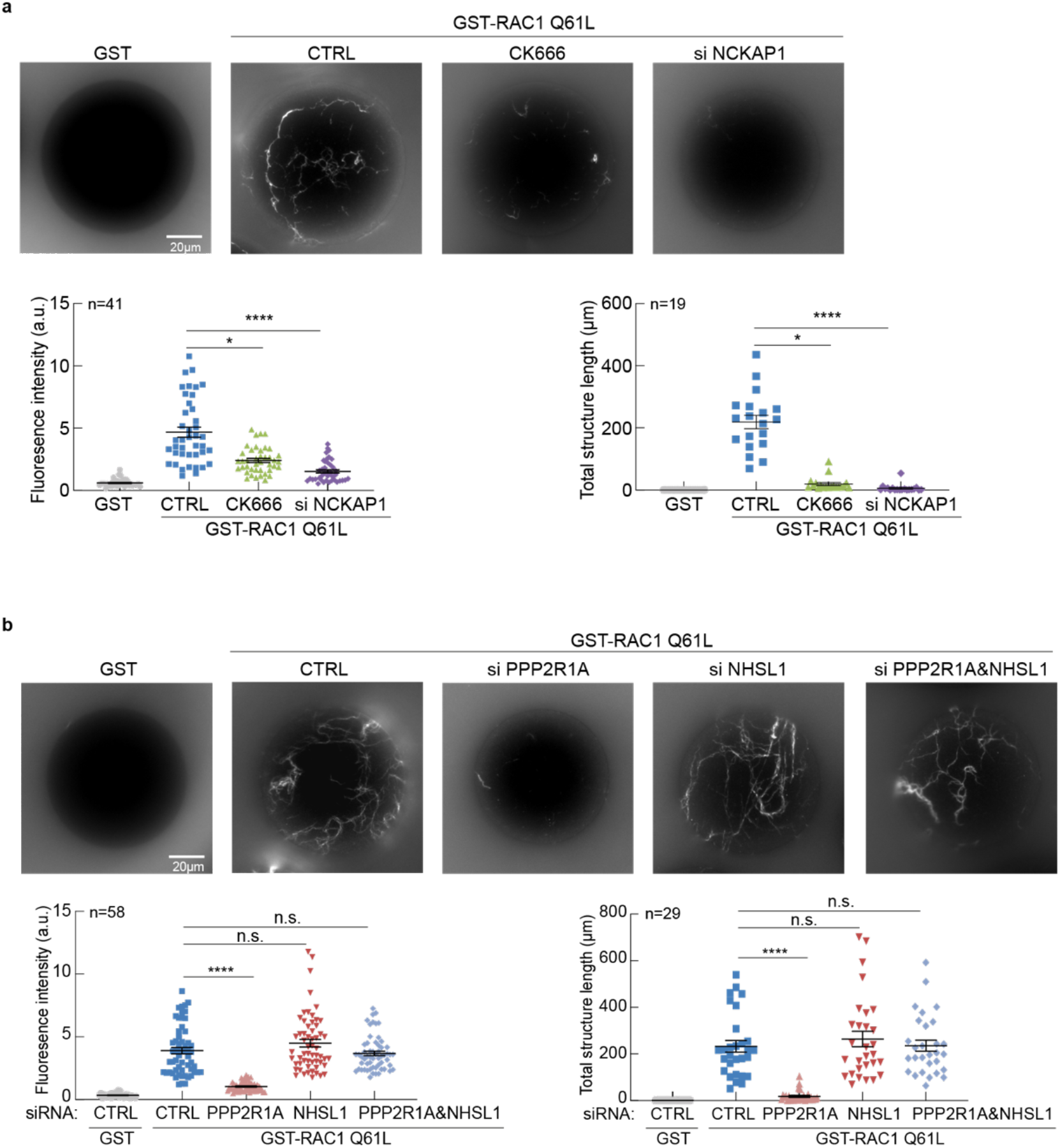
PPP2R1A is required for RAC1-induced actin polymerization in cell-free extracts. **(a)**Beads coated with GST or GST-RAC1 Q61L were incubated with cytosolic extracts of MCF10A cells treated with 200 μM CK666 or depleted of NCKAP1. Rhodamine-labeled actin structures polymerized on the beads were examined by fluorescent microscopy and quantified by average florescence intensity and total structure length. **(b)** Similar experiment performed with cytosolic extracts of MCF10A cells depleted of PPP2R1A, NHSL1 or both. 3 biological repeats with similar results for both experiments. n corresponds to the number of beads used for quantification from all 3 experiments.

### Uncoupling PPP2R1A From WSC Impairs Migration Persistence

To uncouple PPP2R1A from WSC, we mapped PPP2R1A binding sites in NHSL1. NHSL1 was divided in four fragments excluding the WHD. Upon transient transfection of 293T cells with GFP fusion proteins of NHSL1 fragments, we found that the 4^th^ fragment at the C-terminus of NHSL1 retrieved a large amount of PPP2R1A upon GFP immunoprecipitation (Fig.6a). The 2^nd^ fragment of NHSL1 that contains the two previously reported binding sites for the SH3 domain of ABI1 ^16^ also retrieved PPP2R1A, but to a much lesser extent than the 4^th^ fragment. The AlphaFold2 software predicted with high confidence three motifs in fragment 4 that interact with the PPP2R1A scaffold or the PPP2R5D regulatory subunit retrieved in the WSC TAP (Fig.6b). The first two motifs recognize the first HEAT repeat of PPP2R1A, but with different binding and probably mutually exclusive modes. The third motif of NHSL1 fragment 4 corresponds to a previously described consensus motif for binding regulatory subunits of the B56 family, such as PPP2R5D ^39^. Upstream of the consensus motif, binding is reinforced by a long stretch of 25 residues containing two successive and conserved proline residues (Fig.S5). We isolated MCF10A cells that stably express NHSL1 fragment 4. Migration persistence was dramatically decreased in these cells compared to controls (Fig.6c, Fig.S5, Movie S9), suggesting that coupling PPP2R1A to the WSC is critical to regulate cell migration. To obtain further evidence of this point, we investigated the activity of mutated PPP2R1A.

**Figure 6.**
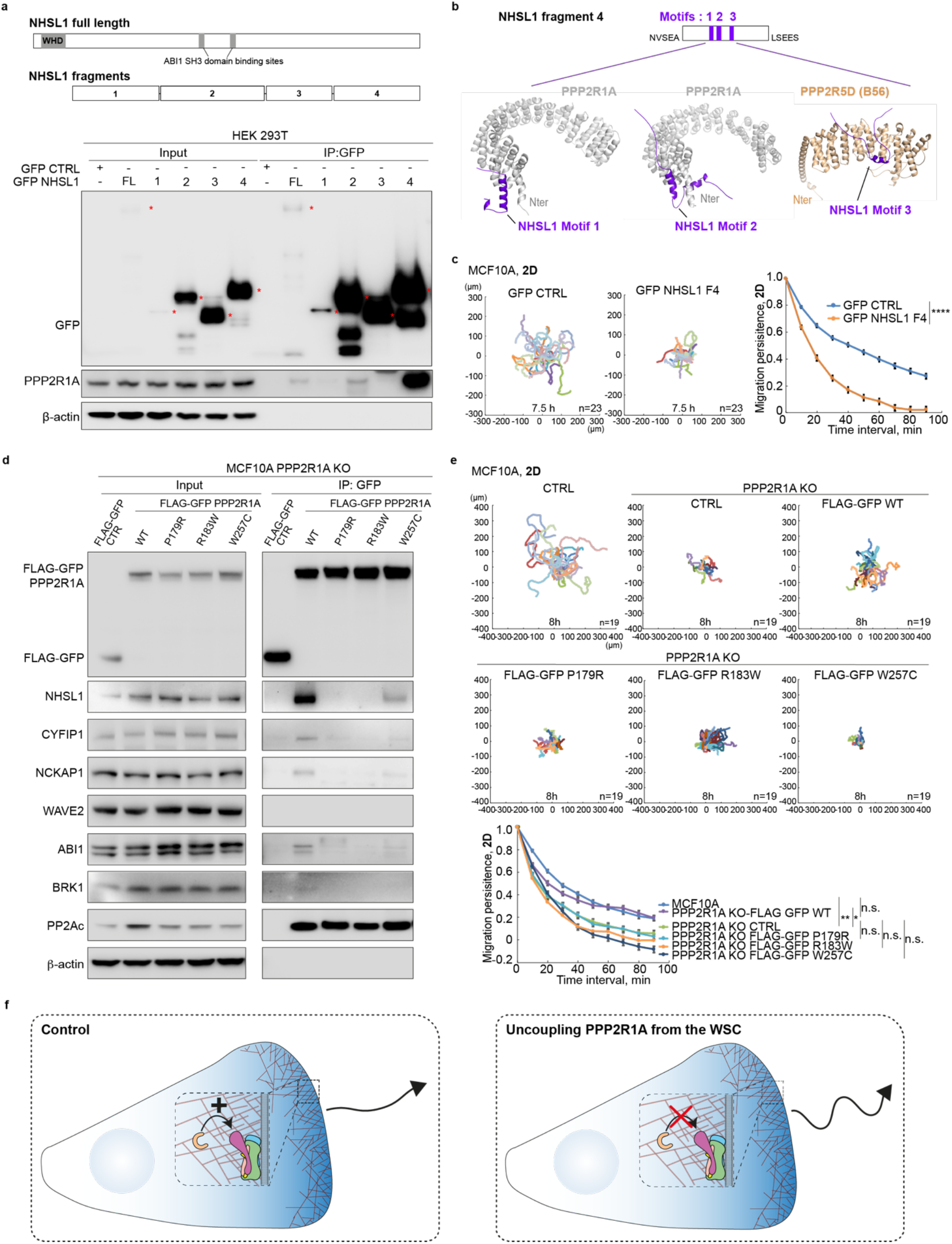
Uncoupling PPP2R1A from the WSC prevents PPP2R1A from regulating migration persistence. **(a)** Scheme of NHSL1 FL and fragments. WHD: WAVE homology domain. HEK 293T cells were transiently transfected with GFP-NHSL1 FL or NHSL1 fragments. Transfected cells were subjected to GFP immunoprecipitation. Western blots with the indicated antibodies. 3 biological repeats of the same experiment with similar results, only one is displayed. (b) Prediction of 3 structural motifs that recognize the scaffolding subunit PPP2R1A or the regulatory subunit PPP2R5D of the PP2A complex using Alphafold2. Cartoon representations of the complexes between PPP2R1A (grey) and motif 1(P_1380_SRP-DDH_1410_ of NHSL1 in violet or motif 2 (A_1430_SP-EPS_1490_ of NHSL1 in violet). Motif 3 (S_1522_LS-EPV_1569_ of NHSL1 in violet) interacts with PPP2R5D (wheat). **(c)** Cell trajectories, migration persistence, cell speed and MSD were extracted from 2D random migration of MCF10A cells stably expressing GFP-NHSL1 fragment 4, n=23. 2 biological repeats of the same experiment with similar results, only one is displayed. **(d)** *PPP2R1A* knockout MCF10A cells were isolated. The KO line was stably transfected with plasmids expressing GFP-tagged WT or mutant forms of PPP2R1A from tumors of patients. Extracts were subjected to GFP immunoprecipitation and immunoprecipitates were analyzed by Western blots with antibodies that recognize WRC, WSC and the catalytic subunit of the PP2A phosphatase complex. 3 biological repeats with similar results. **(e)** 2D cell trajectories and migration persistence of MCF10A parental cells, KO and rescue derivatives. 3 biological repeats with similar results, only one is displayed. n=19. **(f)** Model showing that PPP2R1A can be uncoupled from the WSC at the lamellipodial edge using the fragment 4 of NHSL1 or tumor associated mutations. *P<0.05; **P<0.01; ****P<0.0001.

*PPP2R1A* is frequently mutated in tumors, especially in gynecologic cancers ^40^. We first obtained a *PPP2R1A* KO MCF10A cell line by transfecting purified Cas9 and gRNA, and established that both *PPP2R1A* alleles were inactivated by deletions creating a frameshift (Fig.S6). We then isolated stable transfectants from this KO expressing FLAG-GFP PPP2R1A fusion proteins, either wild type or harboring the frequent cancer-associated mutations, P179R, R183W or W257C. GFP immunoprecipitations showed that the three mutations strongly impair binding to the WSC, but not to the PP2A catalytic subunit (Fig.6d). In the 2D cell migration assay, *PPP2R1A* KO MCF10A cells displayed reduced migration persistence, like PPP2R1A knocked-down cells. Whereas wild type PPP2R1A rescued the phenotype, mutated forms of PPP2R1A were unable to restore persistence (Fig.6e, Movie S10). This experiment confirmed that the interaction of PPP2R1A with the WSC is important for migration persistence.

We then differentiated acini from the stable rescued cell lines at the surface of Matrigel. *PPP2R1A* KO MCF10A cells were impaired in their ability to differentiate hollow acini with correct cell polarity, a hallmark of cell transformation (Fig.S6). The multicellular structures formed instead were also less regularly spherical than controls. These phenotypes were rescued by wild type PPP2R1A, but not with the derivatives harboring a mutation. These results confirmed that the tumor-associated mutations are loss-of-function and suggested that the interaction of PPP2R1A with the WSC is important to prevent cancer progression.

## DISCUSSION

In this study, we uncovered a novel regulator of migration persistence, PPP2R1A, using differential proteomics when RAC1 was activated and when the formation of its downstream product, branched actin, was impaired. PPP2R1A is a well-characterized scaffold subunit of the PP2A complex, an abundant and heterogeneous phosphatase that regulates several signaling pathways ^41^. PP2A phosphatase activity has previously been connected to cell migration and tumor cell invasion ^42–44^, but there is no evidence that the precise role of PPP2R1A we report here in regulating migration persistence depends on PP2A phosphatase activity. Indeed, the phosphorylated sites that we identified on WSC or WRC subunits did not depend on the presence of PPP2R1A. PPP2R1A may ‘moonlight’ and regulate migration persistence independently of PP2A phosphatase activity. Yet there is no mutual exclusivity of PPP2R1A interaction with the WSC and assembly of PP2A complexes, since we detected PP2A catalytic and regulatory subunits when we selected the WSC by TAP. The amounts of PP2A catalytic subunits retrieved by the TAP of ABI1 upon RAC1 activation or inhibition of branched actin did not show the same variations as that of PPP2R1A, suggesting different pools of PPP2R1A with varying degrees of PP2A assembly. In line with this idea, a large-scale proteomics-based quantification of protein abundance has previously revealed that PPP2R1A was almost one order of magnitude more abundant than PP2A catalytic and regulatory subunits with which it associates ^45^.

The A-type subunit PPP2R1A is composed of 15 HEAT repeats, onto which the catalytic C subunit and one of the various B regulatory subunits assemble to form the PP2A holoenzyme ^41^. PPP2R1A is mutated in various cancers, particularly often in endometrial cancers (up to 20 % in some subtypes) ^46^. PPP2R1A is considered a tumor suppressor but has atypical characteristics. Tumor-associated PPP2R1A substitutions fall into hotspots, like mutations activating oncogenes. Most point mutations cluster in HEAT repeats 5 and 7, where they have been shown to impair the interaction with B regulatory subunits to a varying extent depending on the mutation and the B subunit considered ^47,48^. Only one of the two *PPP2R1A* alleles is mutated in tumors, which is also classical of oncogenes. Haploinsufficiency and dominant negative effects are thought to account for PPP2R1A inactivation ^47,49^. Here, using a *PPP2R1A* knock-out cell line and examining the rescue provided by wild type or mutated forms (P179R and R183W in HEAT repeat 5 and W257C in HEAT repeat 7), we unambiguously showed that these tumor-associated mutations inactivate PPP2R1A with respect to this new function in migration persistence as well as in the assay of differentiating acini in Matrigel. Inactivating mutations are consistent with the tumor suppressor nature of PPP2R1A and haploinsufficiency.

Tumor-associated mutations of PPP2R1A impair binding to the here uncovered WSC, where NHSL1 replaces the WAVE subunit. This alternative form of WAVE complex was suspected for a long time, based on the presence of a WAVE homology domain in Nance-Horan Syndrome family proteins ^19^, but was not previously reported. NHSL1 also associates with WRC through two binding sites for the SH3 domain of ABI1 ^16^. It is probably because NHSL1 exists in different pools and complexes, as does PPP2R1A, that these two proteins can regulate migration persistence in opposite ways, positive for PPP2R1A and negative for NHSL1. PPP2R1A, however, selects a specific conformation of NHSL1, where this largely unstructured protein occupies the “shell” composed of all WRC subunits but WAVE. NHSL1 interactions within WSC appear less extensive than WAVE interactions within WRC. NHSL1 is thus unlikely to chase WAVE away but would rather occupy its position when WAVE is not there.

The WSC likely corresponds to a novel intermediate in the WRC life cycle, possibly as a way of activating WAVE. We have obtained evidence that PPP2R1A is required for polymerization of branched actin in cell extracts and for persistence in random and directed migration assays. The surprise is that this requirement of PPP2R1A is not compulsory and depends on the presence of NHSL1 in all assays. The PPP2R1A-WSC interaction may contribute to WRC activation in addition to the primary mechanism, which has been reconstituted *in vitro* without PPP2R1A or NHSL1: Active RAC1 activates the WRC through a conformational change that exposes the Arp2/3 activating WCA domain of WAVE ^6,7^. Even if NHSL1 cannot chase WAVE away, the existence of the WSC suggests that the WAVE subunit can dissociate from the WRC that maintains it inactive ^4,5^. Freeing WAVE from WRC would be another way to expose its Arp2/3 activating WCA motif. It is difficult to anticipate the mechanism by which WAVE would disengage from the WRC. WAVE proteins were reported to be degraded by proteasomes upon cell activation and ubiquitylated on a specific lysine residue in their WHD ^50,51^. But these reports do not demonstrate that this is a free form of WAVE that is degraded after activation. There are many possible views of the WSC role in the WAVE life cycle. The in vitro reconstitution assay of RAC1-dependent branched actin structures we report here using cell extracts is likely to play an important role in testing different scenarios for a multistep WAVE activation cycle.

Migration persistence is thought to depend on positive feedback ^20^, which is manifested in propagating waves of branched actin polymerization ^22^. Positive feedback has been observed at multiple molecular levels. The simple fact that the product of the Arp2/3 reaction is a new actin filament that can become the substrate of a new Arp2/3-dependent branching reaction is a positive feedback referred to as an autocatalytic reaction ^52^. The WRC turns over at the lamellipodium edge due to elongation of actin filaments that it contributes to generating ^24,53^. Coronin1A decorates lamellipodial actin and further activates RAC1 via its association with ArhGEF7 ^54^. Similarly, we found PPP2R1A in the width of the lamellipodium, ideally localized to sense lamellipodial actin, whereas PPP2R1A association with the WSC is likely to take place at the lamellipodial edge, where NHSL1 is localized, and where new actin filaments are nucleated by the Arp2/3 (Fig.6f). This does not fully demonstrate involvement in feedback, but we initially focused on PPP2R1A, among the many ABI1 partners, because its interaction with the WSC was regulated by branched actin. It is worth noting in this respect that simulations using molecular dynamics have revealed that the HEAT-repeat scaffold of PPP2R1A is particularly elastic and might play a role in mechanotransduction ^55^. Furthermore, PPP2R1A is dispensable for migration persistence when RAC1 is constitutively activated by the Q61L mutation. All of this evidence points to a possible role for PPP2R1A, and the WSC it associates with, in a positive feedback that sustains directional migration through continuous actin polymerization at the leading edge. Future work should aim to find ways to dissect, at the molecular level, the complex circuitry that mediates feedback and persistence.

## Author Contributions

YW performed most of the experiments and participated in writing of the manuscript. GC performed the mass spectrometry under the direction of JV. SR performed live localizations of fluorescent fusion proteins. RG generated the structural models. MK generated NHSL1 expression plasmids. CD, AB and AIB helped to set up the haptotaxis assay. AMG and AP have jointly supervised the work and wrote the manuscript.

## Acknowledgments

We thank Vassilis Koronakis for guidance in reconstitution assays using cell extracts. We thank Anna Castro, Gregory Giannone and Emmanuel Derivery for critical reading of the manuscript. This work was supported by grants from Agence Nationale de la Recherche (ANR-20-CE13-0016-01) and Fondation ARC pour la Recherche sur le Cancer (ARC PJA 2021 060003815). Mass spectrometry equipment was subsidized by Conseil Régional d’Ile-de-France (Sesame N°10022268). The PhD of YW was supported by fellowships provided by Fondation pour la Recherche Médicale and by Fondation ARC pour la Recherche sur le Cancer.

