## Supplementary Information for "PPP2R1A Regulates Migration Persistence through the WAVE Shell Complex"

**Methods …………………………………..… 2**

**Supplementary Figures ………………...… 10**

**Movie Legends ……………………………. 18**

**Supplementary References ………………. 20**

#### METHODS

##### Cells

MCF10A cells were grown in DMEM/F12 medium supplemented with 5% horse serum, 20 ng/mL epidermal growth factor, 10µg/mL insulin, 100 ng/mL cholera toxin, 500 ng/mL hydrocortisone and 100U/mL penicillin/streptomycin. MDA-MB-231 and B16-F1 cells were grown in DMEM medium with 10% FBS and 100 U/mL penicillin/streptomycin. Medium and supplements were from Life Technologies and Sigma. Cells were incubated at 37°C in 5% CO2.

##### Plasmids

The following plasmids were built using a building block method [1]:
MXS AAVS1L SA2A Puro bGHpA EF1FLAG GFP Blue2 Sv40pA AAVS1R (control)
MXS AAVS1L SA2A Puro bGHpA EF1 FLAG GFP ABI1 Sv40pA AAVS1R
MXS AAVS1L SA2A Puro bGHpA EF1 FLAG GFP PPP2R1A Sv40pA AAVS1R
MXS PGK Blasti bGHpA CAG PC HA Blue2 Sv40pA (control)
MXS PGK Blasti bGHpA CAG PC HA BRK1 Sv40pA
MXS PGK ZeoM bGHpA EF1Flag mScarlet PPP2R1A SV40pA
The plasmid pCAG-EGFP-NHSL1-IRES-Puro encoding GFP fusion with FL NHSL1 containing the WHD (1-1639). FL NHSL1 corresponds to the isoform X6 (XP_047275069.1) with 7 substitutions T440I, A449V, T891A, G1082E, A1123T, P1329S and S1466N. The fragments 1 (MVVFI…KSRDH), 2 (LISRH…EGSGT), 3 (MKKLD…VEPAE) and 4 (NVSEA…LSEES) correspond to the isoform X19 (XP_047275073.1) with 7 substitutions T363I, A372V, T814A, G1005E, A1046T, P1252S and S1389N and were expressed as GFP fusion proteins from the same plasmid as the full-length NHSL1.

##### Stable Cell Lines

Stable transfections of MCF10A and MDA-MB-231 cells were performed using Lipofectamine 3000 (Invitrogen) with the plasmids encoding FLAG-GFP, FLAG-GFP ABI1, FLAG-GFP PPP2R1A, FLAG-GFP NHSL1 Fragment 4. To obtain stable integration at the AAVS1 site, cells were co-transfected with two TALEN constructs (Addgene #59025 and 59026) inducing a double strand break at the AAVS1 locus [2]. Cells were selected with 1µg/mL puromycin (Invivogen) and pooled.

Stable MCF10A double cell line was obtained by transfecting cells stably expressing FLAG-GFP PPP2R1A with PC-HA BRK1 or PC-HA Blue2. Cells were then selected with 10µg/ml of Blasticidin (Invivogen). Single clones were expanded and analyzed by Western blot.

**Knockdown and Knockout**

MCF10A and MDA-MB-231 knockdown cells were obtained by transfecting 20 nM siRNAs (Dharmacon ON-TARGET SMART Pool: L-010259-00-0010 for PPP2R1A, L-032698-00-0010 for NHSL1) with Lipofectamine RNAiMAX (Invitrogen). After 3 days, cells were subjected to videomicroscopy or Western blot.

MCF10A knockout cell lines were generated with CRISPR/Cas9 system. The following gRNAs were used:

*PPP2R1A* 5'-CATAGACGAACTCCGCAATG-3'

*NHSL1* 5'-TCGGCTTTCCTCATCTAGGT-3'

non-targeting 5'-AAAUGUGAGAUCAGAGUAAU-3'.

Cells were transfected with gRNA, tracrRNA and Cas9 protein by Lipofectamine CRISPRMAX™ (all reagents from ThermoFischer Scientific). After 2 days, cells were diluted at 0.8 cells/well in 96-well plates. Single clones were expanded and analyzed by Western blot. The positive clones were confirmed by sequencing.

**Antibodies**

The antibodies used were: anti-PPP2R1A (Bethyl Laboratories, A300-962A); anti-NHSL1 (Sigma-Aldrich, HPA029967); anti-PP2Ac (Bethyl Laboratories, A300-732A); anti-GFP (Roche, 11814460001); anti-NCKAP1 (Bethyl Laboratories, A305-178A); anti-WAVE1 (R&D Systems, AF5514); anti-WAVE3 (R&D Systems, AF5515); anti-phospho-WAVE2 Ser308 (Millipore, 07-1511); anti-GAPDH (Thermo Fisher Scientific, AM4300); anti-β-actin (Thermo Fisher Scientific, AM4302). Home-made CYFIP1, ABI1, WAVE2 antibodies and BRK1 antibody were described previously [3,4].

**Western Blots**

Cells were lysed in XB-NP40 buffer (50mM HEPES, 50mM KCl, 1%NP-40, 10mM EDTA, pH 7.7) supplemented with protease inhibitor (Roche), the lysates were clarified by centrifugation at 13,000 rpm for 15 min and subjected to SDS-PAGE using NuPAGE 4-12% Bis-Tris or 3-8% Tris-Acetate gels (Life Technologies). Nitrocellulose membranes were incubated with primary antibodies, HRP conjugated secondary antibodies (Sigma) and developed with SuperSignal™ West Femto Substrate (Thermo Fisher Scientific) and ChemiDoc imaging system (BIO-RAD). Densitometry of Western blots was performed with ImageJ.

**Immunoprecipitation and Tandem Affinity Purification**

Cells stably expressing FLAG-GFP ABI1 or FLAG-GFP PPP2R1A were lysed with XB-NP40 buffer (50mM HEPES, 50mM KCl, 1%NP-40, 10mM EDTA, pH 7.7) supplemented with protease inhibitors at 4°C for 30 min. The phosphatase inhibitor cocktail PhosSTOP (Roche) was added for phosphosite analysis. The lysates were clarified by centrifugation at 13,000 rpm for 15 min. Clarified cell extracts were incubated with FLAG-M2 beads (Sigma) at 4°C for 4 h. FLAG-M2 beads were washed with XB-NP40 buffer and eluted with 0.5 mg/ml FLAG peptide (Sigma) in XB (50mM HEPES, 50mM KCl, 10mM EDTA, pH 7.7) overnight at 4°C. FLAG elutions were collected and incubated with GFP-trap beads (Chromotek) at 4°C for 1 h. The GFP-trap beads were washed with XB-NP40 buffer. 20% of the beads were subjected to SDS-PAGE for Western blot or silver staining (SilverQuest™ Silver Staining Kit, Thermo Fisher Scientific). 80% of the beads were analyzed by mass spectrometry.

To purify the WSC, MCF10A cells stably expressing FLAG-GFP PPP2R1A and PC-HA BRK1 were lysed with XB-NP40 buffer (50mM HEPES, 50mM KCl, 1%NP-40, 10mM EDTA, pH 7.7) supplemented with protease inhibitors at 4°C for 1 h, then the lysates were clarified by centrifugation at 13,000 rpm for 15 min. Cell extracts were incubated with FLAG-M2 beads (Sigma) overnight at 4°C. FLAG-M2 beads were washed with XB-NP40 buffer, and eluted with 0.5 mg/ml FLAG peptide (Sigma) in FLAG-elution buffer (50mM HEPES, 50mM KCl, 1mM $CaCl_2$, pH 7.7). FLAG elutions were incubated with PC beads (Anti-Protein C Affinity Matrix, Sigma) overnight at 4°C. PC beads were washed eluted with EGTA-elution buffer (50mM HEPES, 50mM KCl, 10mM EGTA, pH 7.7) overnight. PC elutions were subjected to SDS-PAGE and mass spectrometry.

**Mass Spectrometry**

The resins containing the immunoprecipitated sample were loaded onto a 10 kDa cutoff centrifugal filters (Microcon, Millipore-Merck) and washed with 500 µL of ammonium bicarbonate buffer (50 mM, pH 8.0, AMBIC). Disulfide reduction was performed adding to the centrifugal filters 200 µL of a solution containing 10 mM dithiothreitol in AMBIC for 2h at 37 °C. Thiol alkylation was performed by adding to the previous samples 200 µL of a solution containing 50 mM iodoacetamide in AMBIC for 30 minutes at room temperature. Reagents were removed by filtration and sample washed three times with 500 µL of AMBIC. Proteins were digested with 1 µg of trypsin/Lys-C (Promega) in 100 µL of AMBIC overnight at 37 °C. The resulting peptide mixture was filtered and acidified with trifluoro acetic acid at a final concentration of 0.1%. Technical triplicates were systematically analyzed.

For each fraction, 6 µL of sample was concentrated on a C18 cartridge (Dionex Acclaim PepMap100, 5 µm, 300 µm i.d. x 5 mm) and eluted on a capillary reverse-phase column (C18 Dionex Acclaim PepMap100, 3 µm, 75 µm i.d. x 50 cm) at 220 nL/min, with a gradient of 2% to 38% of buffer B in 60 min (buffer A: 0.1% aq. Formic Acid/Acetonitrile 98:2 (v/v); buffer B: 0.1% aq. Formic Acid/Acetonitrile 10:90 (v/v)), coupled with a quadrupole-Orbitrap mass spectrometer (Q Exactive HF, ThermoFisher Scientific) using a Top 20 data-dependent acquisition MS experiment: 1 survey MS scan (400-2,000 m/z; resolution 70,000) followed by 20 MS/MS scans on the 20 most intense precursors (dynamic exclusion of 30 s, resolution 17,500).

Protein identification was performed with MaxQuant search engine v.1.5.3.30 against the human Swiss-Prot database (updated in 07/2020), with the following parameter: methionine oxidation, cysteine carbamidomethylation, asparagine/glutamine deamidation and serine/threonine/tyrosine phosphorylation as variable modifications, first search error tolerance 20 ppm, main error tolerance 6 ppm, MS/MS error tolerance 20 ppm, FDR 1%. Quantification was performed in label-free LFQ normalization mode, using at least 2 Razor or unique peptide per protein. Quantities were estimated using LFQ intensities. Significant changes in protein amounts were estimated by ANOVA with Bonferroni's Post-Hoc test using a p-value cutoff threshold of 0.05.

Raw files of the LC-MSMS analyses and the database researches have been deposited in PRIDE (https://www.ebi.ac.uk/pride/) with the accession number PXD031584. Files with the reference number 170414 refer to the TAP purification of FLAG-GFP-ABI1, 181220 to the TAP purification of FLAG-GFP-PPP2R1A, 210415 to the TAP purification of the WSC and 200120 to the identification of phosphosites in the TAP purification of FLAG-GFP-ABI1. These files can be downloaded using the following username:, and password: DsEdCLt8.

**Migration and Videomicroscopy**

Random cell migration assays were performed in µ-Slide 8 well ibidi dishes. For 2D migration assays, cells were seeded on the dishes coated with 20µg/ml Fibronectin (Sigma). For 3D migration assays, cells were sandwiched between two layers of 2mg/ml collagen (rat tail

collagen type I, Corning). After seeding cells for 24h, videomicroscopy was performed on an inverted Axio Observer microscope (Zeiss) equipped with a Pecon Zeiss incubator XL multi S1 RED LS (Heating Unit XL S, Temp module, CO2 module, Heating Insert PS and CO2 cover), a definite focus module and a Hamamatsu camera C10600 Orca-R2. Images were acquired every 5 or 10 min for 24 h with 10x objective for 2D migration, and every 10 min for 48 h with 20x objective for 3D migration. Individual cells were tracked by ImageJ software-Manual Tracking plug-in. DiPer software was used to analyze the cell migration parameters [5].
Fibronectin gradients were prepared with PRIMO photopatterning system (Alvéole). 35mm ibidi dishes with glass bottom were treated with plasma for 1 min. PDMS stencils (Alvéole) with three 3x3mm wells were stacked on each plasma-treated dish immediately. PDMS stencil wells were coated with PLL-g-PEG for 1 h and rinsed 3 times with PBS. Then photoinitiator (PLPP) (Alvéole) was added to the PDMS stencil wells for micropatterning. LEONARDO photopatterning software was used to design the micropatterns (width 645μm, height 1031μm with 100% to 0% grayscale gradient). The dishes with PDMS stencils were placed onto the microscope holder (Nikon ECLIPSE Ti2) with PRIMO module, and the patterns were projected to the surface at a UV dose of 1500 mg/mm$^2$. PDMS stencil wells were rinsed 3 times with PBS and coated with 50μg/ml Fibronectin/Fibrinogen-Alexa647 (Invitrogen) for 30 minutes at 37°C. The Fibronectin/Fibrinogen-Alexa647 were only adsorbed on the previously illuminated areas. After rinsing the wells 3 times with PBS, MCF10A cells were seeded in the coated PDMS stencil wells. Cells were washed once with medium after adhering for 2 h. After 24 h, videomicroscopy was performed on an inverted Axio Observer microscope (Zeiss) with the 10x objective. Images were acquired every 10 min for 24 h. Individual cells were tracked by ImageJ software-Manual Tracking plug-in. The tracks obtained were analyzed by the chemotaxis tool (Ibidi) to extract the FMI values and cell trajectory plots. The FMI values were plotted by GraphPad Prism software as mean ± 95% confidence intervals.
B16-F1 cells were transiently transfected with GFP and mScarlet plasmids and analyzed by videomicroscopy after 2 days. Videos were acquired using a confocal laser scanning microscope (TCS SP8, Leica) equipped with a high NA oil immersion objective (HC PL APO 63×/ 1.40, Leica), a white light laser (WLL, Leica) and controlled by the LasX software. Images were taken every 10 seconds for about 5-10 min. Kymographs were drawn using Multi Kymograph tool in ImageJ. To analyze the localization of proteins, radial line scans were performed and analyzed as described [6].

**3D Acini**

Single cells were seeded on a 1 mm thick solidified layer of Matrigel (growth factor reduced, Thermo Fisher Scientific, #CB-40230C) in the 8-well glass chamber (Merck, #PEZGS0816) and grown for 3 weeks in MCF10A medium with 1% horse serum, 5 ng/mL EGF and 2% Matrigel. Then the acini were fixed and stained with indicated antibodies. Images were acquired using a confocal laser scanning microscope (TCS SP8, Leica) equipped with a high NA oil immersion objective (HC PL APO 63×/ 1.40, Leica), a white light laser (WLL, Leica) and controlled by the LasX software.

**Protein Purification**

GST-fusion RAC1 WT, RAC1 Q61L and CDC42 Q61L were expressed in and purified from *E. coli* BL21. After 3 hours induction at 37°C with 1mM IPTG, cells were resuspended and lysed in lysis buffer (50mM Tris-HCl, 100mM NaCl, 2.5mM $CaCl_2$, 10mM $MgCl_2$, 1%Triton X100, 5% glycerol, 0.5 mg/ml lysozyme, 10μg/ml DNAseI, 1mM DTT, pH8.0, protease inhibitors) at 4°C for 1 h. The lysates were clarified by centrifugation at 13,000 rpm for 30 min. The supernatants were incubated with Glutathione Sepharose beads (GE healthcare) at 4°C for 3 h. The beads were washed with TBS buffer (50mM Tris, 100mM NaCl, 2.5mM $CaCl_2$, 5mM $MgCl_2$, 1mM DTT, pH8.0), then the bounded proteins were eluted with GST elution buffer (50mM Tris-HCl, 5mM $MgCl_2$, 10mM glutathione, pH8.0). The elutions were dialyzed in buffer (50mM Tris, 5mM $MgCl_2$, 20% glycerol, pH8.0) and kept at -80°C until use.

**GST Pull-Down**

MCF10A cell extracts were prepared with XB-NP40 buffer as described previously. 20μg purified GST-fusion proteins were incubated with 20μl Glutathione Sepharose beads (GE healthcare) in 500μl incubation buffer (50mM Tris-HCl, 100mM NaCl, 2.5mM $CaCl_2$, 10mM $MgCl_2$, 1%Triton X100, 5% glycerol, 1mM DTT, pH8.0) at 4°C for 1 h. The pre-coated beads were washed and incubated with 1 ml MCF10A cell extract at 4°C for 1 h. The beads were washed with XB-NP40 buffer and subjected to Western blot.

**Actin Polymerization in Cell-Free Extracts**

MCF10A cells were lysed either by nitrogen cavitation (Parr instruments, 500 Psi for 20 minutes) in buffer (50mM HEPES, 50mM NaCl, 5mM $MgCl_2$, 0.1mM EDTA, 1mM DTT, pH 7.7, protease inhibitor), or with NP40 containing buffer (50mM HEPES, 50mM KCl, 5mM $MgCl_2$, 1%NP-40, pH 7.7, protease inhibitor) at 4°C for 30 min. The NP40 containing extract supported RAC1 Q61L-induced actin polymerization, whereas extract obtained by nitrogen cavitation supported CDC42 Q61L-induced actin polymerization. The extracts were clarified by centrifugation at 13,000 rpm for 15 min. 10µl clarified cell extract was supplemented with 2µl energy mix (20mM ATP, 150mM creatine phosphate, 20mM $MgCl_2$, 2mM EGTA) and 0.75µl rhodamine-actin (1mg/ml, Cytoskeleton, Inc.). The mixture was centrifuged 5 min at 13,000 rpm. To trigger the reaction, 1µl Glutathione Sepharose beads bound with 2µg GST-RAC1 Q61L or GST-CDC42 Q61L were added to 10µl reaction mix. After 1 h incubation at room temperature, the reaction was squashed in between a coverslip and the microscope slide. The coverslip was sealed with melted VALAP (Vaseline-Lanoline-Paraffin). The beads were observed under an inverted microscope (Olympus IX83) with 60x oil objective. Fluorescence intensity and structure length on the surface of the beads were measured using ImageJ.

**Structural Modeling**

Sequences of human CYFIP1, NCKAP1, BRK1, ABI2, PPP2R1A and PPP2R5D were retrieved from UniProt database [7] and the full-length NHSL1 cloned in plasmid pCAG were used as input of mmseqs2 homology search program [8] with 3 iterations to generate a multiple sequence alignment (MSA) against the UniRef30 clustered database. Homologs sharing less than 25% sequence identity or less than 50% of coverage of the aligned region with their respective query, were discarded. In case several homologs belonged to the same species, only the one sharing highest sequence identity to the query was kept. Full-length sequences of selected homologs were retrieved and realigned with mafft [9]. To model WSC structure, concatenated MSAs of CYFIP1, NCKAP1, ABI2 (1-160), BRK1 and NHSL1 (1-95, 1-123, or 1-200) were analyzed. Homologs of different subunits belonging to the same species were aligned in a paired manner otherwise in concatenated MSAs. Final concatenated MSAs of WSC contained 2711 positions and 1577 species. MSAs of NHSL1 motif 1 ($P_{1380}$SRP-DD$H_{1410}$), motif 2 ($A_{1430}$SP-EP$S_{1490}$) were similarly concatenated with that of PPP2R1A and MSA of motif 3 ($S_{1522}$LS-EP$V_{1569}$) with that of PPP2R5D (80-530), yielding 3 MSAs from 1733 and 2445 species, respectively. Each concatenated MSA was then used as input to run 5 independent runs of the AlphaFold2 algorithm with 6 iterations each time [10] in order to generate 5 structural models using a local version of the ColabFold interface [11] trained on the multimer dataset [12] on a local HPC equipped with NVIDIA Ampere A100 80Go GPU cards. Best models of each of the 5 runs converged toward similar conformations for each of the 4 modeled molecular systems. High-confidence quality scores of pLDDT in the range of [84.6, 86.2], [91.8, 93.1], [88.3, 88.9], [87.2, 88.9] and pTMscore in the range [0.807, 0.833], [0.816, 0.842], [0.785, 0.8],

[0.844, 0.854] were obtained for WSC and the complexes involving NHSL1 motifs 1, 2 and 3, respectively. For each of the four models, the models with highest pTMscores were relaxed using Rosetta relax protocols to remove steric clashes [13] with strong backbone constraints (std dev. of 0.5 Å for atomic positions) and were used for structural analysis.

The models of i) the Wave Shell Complex (WSC) composed of NHSL1(1-95), CYFIP1, NCKAP1, BRK1 and ABI2(1-160), ii) the NHSL1(1382-1410)-PPP2R1A complex, iii) the NHSL1(1430-1490)-PPP2R1A complex and iv) NHSL1(1522-1569)-PPP2R5D(80-530) complex, are available in ModelArchive (modelarchive.org) with the accession codes ma-agzek (passwd: nFB3QBkRYT), ma-ne9d4 (passwd: KJO5ik7Ksw), ma-sx8ix (passwd: Ozduh9k3eW) and ma-rop1i (passwd: BKiXtzlnG9), respectively.

**Statistics**

Migration persistence for individual cells is evaluated based on the exponential decay and plateau fit as shown below.

$$P = (1 - b) * e^{-\frac{t}{a}} + b$$

Where, $P$ is the migration persistence, $b$ is plateau value, $t$ is the time interval and $a$ is the decay constant. Then the related statistical analysis was conducted through custom-made R programs, as previously described [14].

For other statistical analysis, GraphPad Prism software and Microsoft Excel were used. ANOVA followed by post hoc Tukey's multiple comparison test or Kruskal-Wallis test followed by post hoc Dunn's multiple comparison test were applied based on the Shapiro-Wilk normality test.

Four levels of significance were distinguished: $*P < 0.05$, $**P < 0.01$, $***P < 0.001$, $****P < 0.0001$.

### SUPPLEMENTARY FIGURES

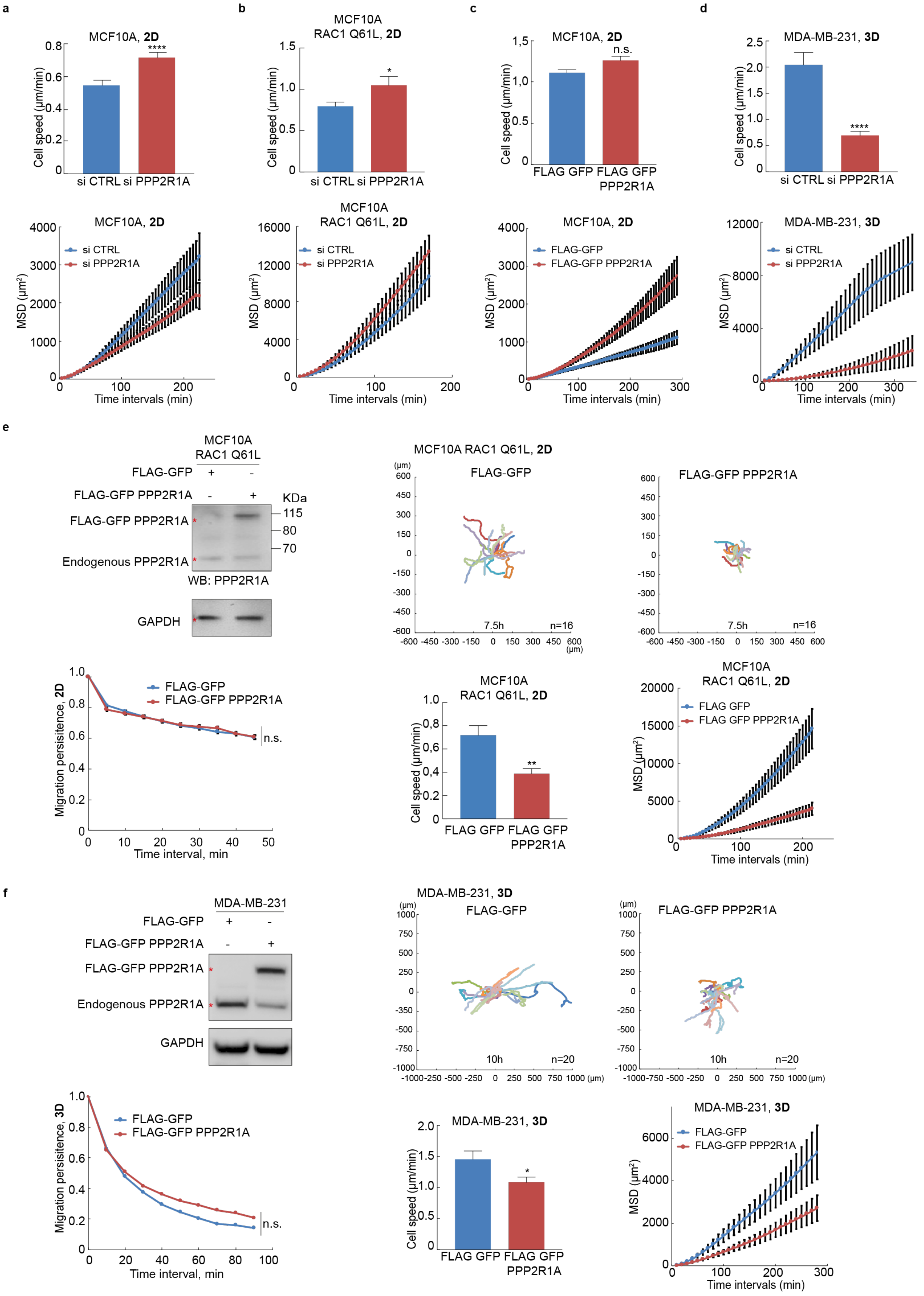

**Figure S1, related to figure 1. Migration parameters of MCF10A and MDA-MB-231. (a-d)** Cell speed and MSD extracted from cell trajectories displayed in corresponding panels of figure 1. 3 biological repeats of the same experiment with similar results, only one is displayed. **(e)** Cell speed and MSD extracted from random migration of single MCF10A cells expressing FLAG-GFP or FLAG-GFP PPP2R1A in 2D, n=16. 2 biological repeats with similar results, only one is displayed. **(f)** MDA-MB-231 cell lines were stably transfected with plasmids expressing FLAG-GFP or FLAG-GFP PPP2R1A and analyzed by Western blots with PPP2R1A or GAPDH antibodies. Cell trajectories, migration persistence, speed and MSD extracted from 3D migration of single MDA-MB-231 cells in collagen type I gels, n=20. 3 biological repeats with similar results, only one is displayed. *P<0.05; ** P<0.01; ****P<0.0001

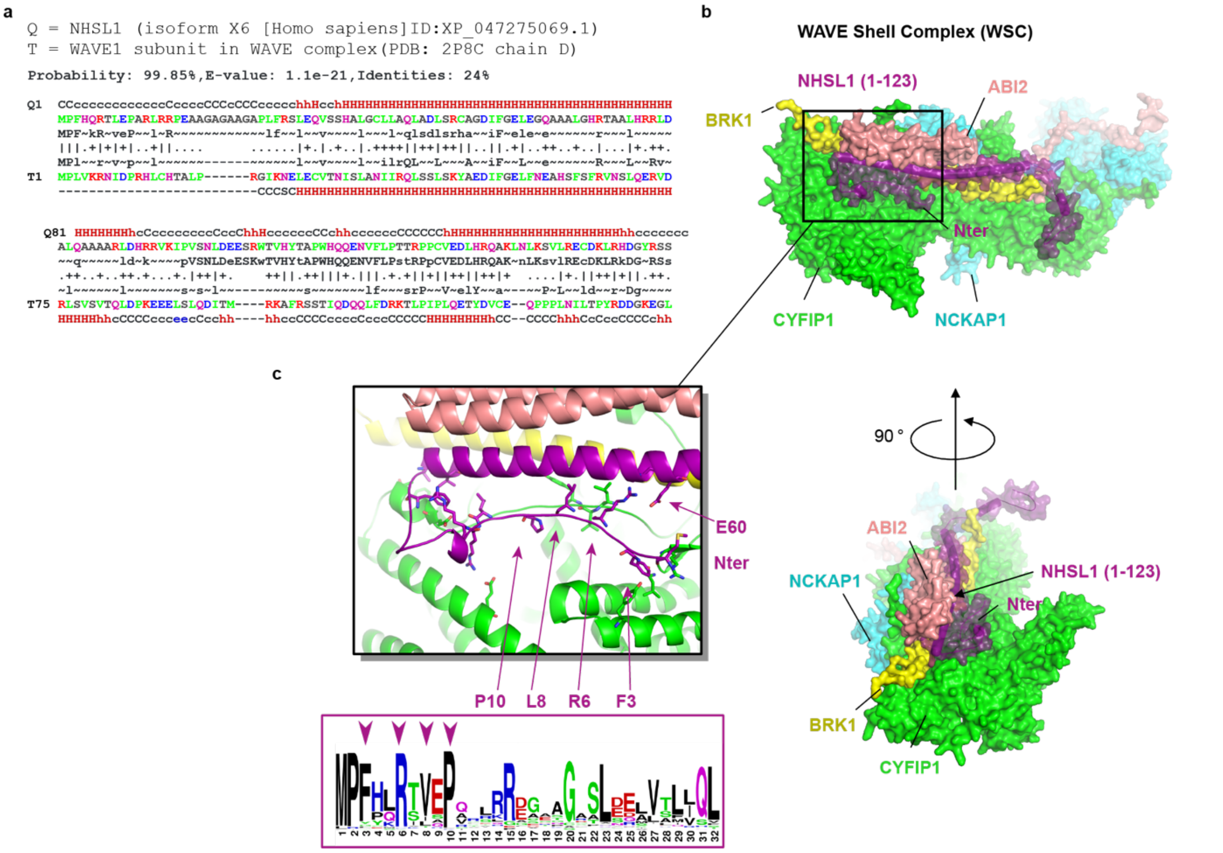


**Figure S2. Structural model of the WSC. (a)** Sequence alignment between the profile of NHSL1 (line Q, 1-160 region) and that of the WAVE1 subunit (line T) extracted from the structure of the WAVE Regulatory Complex (PDB:3P8C). **(b)** Surface representation of the structural model of the WSC generated by AlphaFold2. The model is composed of NHSL1 (1-123) (purple), full-length CYFIP1 (green), NCKAP1 (cyan), BRK1 (yellow) and ABI2 (1-160) (salmon) and is displayed under two views rotated by 90° around the vertical axis. Sensitivity and precision of AlphaFold2 decreases when using long inputs as full WSC. Therefore we also explored potential binary complexes that could be formed between NHSL1 and either CYFIP1 or NCKAP1. These binary structural models with either CYFIP1 or NCKAP1 were very similar to the structural model obtained with full WSC. This suggests that additional interactions downstream of the displayed region of NHSL1 are unlikely. **(c)** Close-up view with cartoon representation. Side chains of amino-acids highlighted as sticks correspond to conserved residues in the N-terminus of NHSL1. Conservation pattern represented in a logo plot using WebLogo[15] indicate the highlighted residues by purple triangles.

a

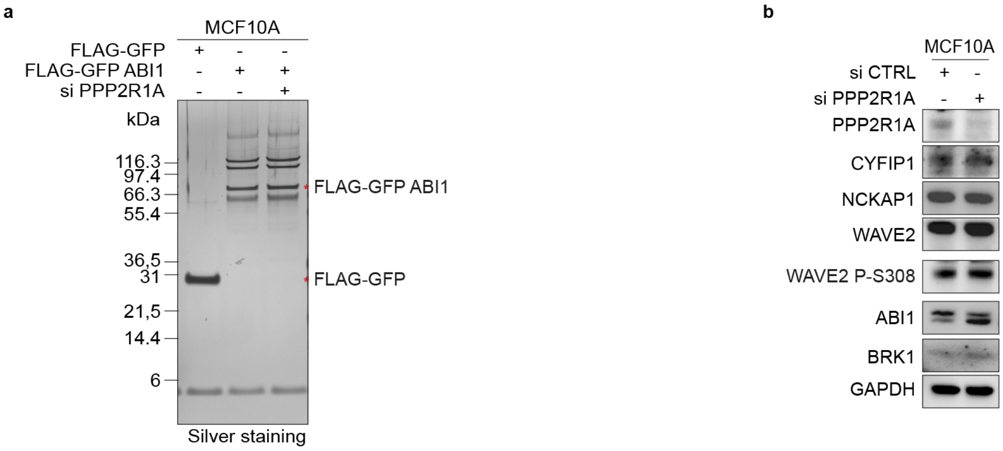


| Protein | Gene names | Number of Phospho (STY) | Position with highest probablity | Probability that the different STY residues are phosphorylated | Ctrl | shPPP2R1A |
|---|---|---|---|---|---|---|
| ABI1 | *ABI1* | 1 | 183 | TNPPTQKPPS(1)PPMSGR | 100 ± 12 | 73 ± 9 |
| ABI1 | *ABI1* | 2 | 183 | T(0.004)NPPT(0.015)QKPPS(0.964)PPMS(0.509)GRGT(0.508)LGR | 100 ± 9 | 0 |
| ABI1 | *ABI1* | 1 | 200 | NT(0.013)PY(0.053)KT(0.934)LEPVKPPTVPNDYMTSPAR | 100 ± 49 | 0 |
| ABI1 | *ABI1* | 2 | 183 | TNPPTQKPPS(0.998)PPMS(0.244)GRGT(0.758)LGR | 100 ± 34 | 0 |
| NHSL1 | *NHSL1* | 1 | 1229 | AVPS(0.986)PT(0.011)T(0.003)GEEGSVHSR | 100 ± 6 | 75 ± 9 |
| NHSL1 | *NHSL1* | 1 | 324 | LDSDAGFHS(1)LPR | 100 ± 47 | 46 ± 0 |
| NHSL1 | *NHSL1* | 2 | 1384 | NHS(0.984)PS(0.988)PPVT(0.029)PTGAAPSLASPK | 100 ± 15 | 105 ± 0 |
| NHSL1 | *NHSL1* | 1 | 1186 | SPGAPSAGEAEARPS(0.001)PS(0.002)T(0.002)T(0.002)PLPDS(0.583)S(0.355)PS(0.054)R | 100 ± 0 | 106 ± 45 |
| NHSL1 | *NHSL1* | 1 | 209 | RKT(0.997)IT(0.003)GVPDNIQK | 100 ± 31 | 64 ± 14 |
| WAVE2 | *WASF2* | 1 | 293 | RS(0.209)S(0.765)VVS(0.026)PSHPPPAPPLGSPPGPK | 100 ± 19 | 53 ± 12 |
| WAVE2 | *WASF2* | 1 | 308 | SSVVS(0.004)PS(0.003)HPPPAPPLGS(0.992)PPGPK | 100 ± 0 | 262 ± 97 |
| WAVE3 | *WASF3* | 1 | 235 | LSQSVYHGAS(0.001)S(0.002)EGS(0.017)LS(0.918)PDT(0.063)R | 100 ± 14 | 57 ± 13 |

**Figure S3. Analysis of phosphorylated sites in WRC and WSC in presence or absence of PP2R1A. (a)** MCF10A stably expressing FLAG-GFP ABI1 were transfected with siRNAs targeting PPP2R1A or control siRNAs. TAP purification of ABI1 was analyzed by SDS-PAGE and by mass spectrometry. Label-free quantification of phosphosites identified by mass spectrometry in relevant subunits of WRC and WSC. 3 technical repeats, mean ± sem. **(b)** Western blots of WRC and WSC subunits including phospho-serine 308 of WAVE2.

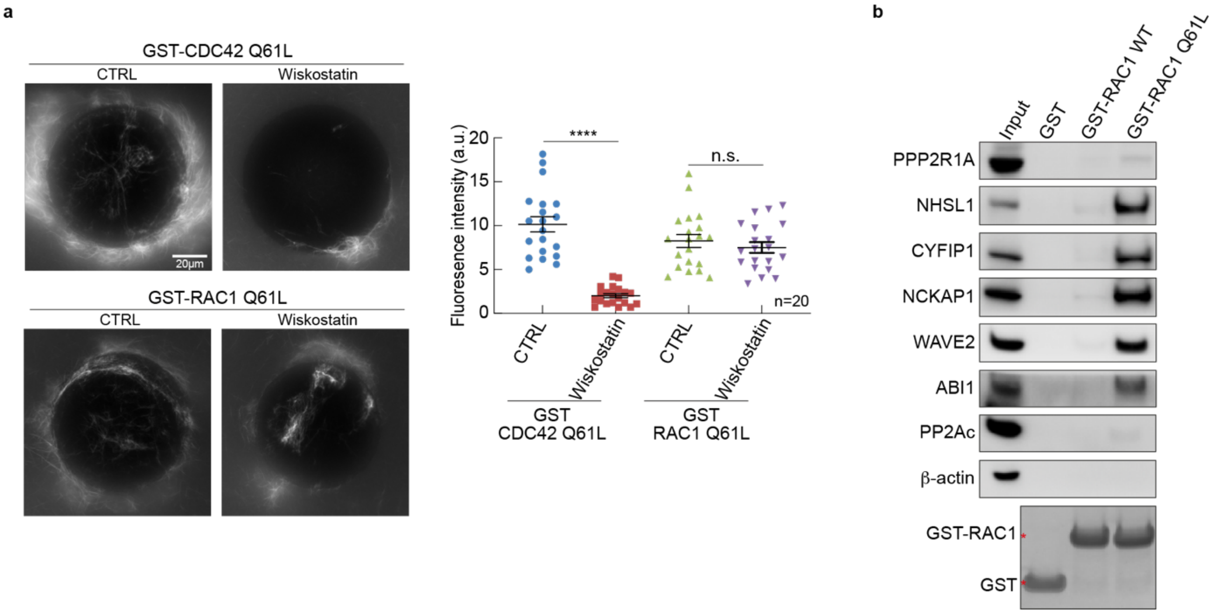


**Figure S4, related to Fig.5. Actin polymerization at the surface of beads displaying GST fusion proteins with small GTPases. (a)** Beads coated with GST-CDC42 Q61L or GST-RAC1 Q61L were incubated with the cell extracts of MCF10A treated or not with 10 µM wiskostatin. Structures containing rhodamine-labeled actin were examined at the surface of beads by epifluorescence and their intensity quantified. **(b)** Beads coated with GST, GST-RAC1 WT or GST-RAC1 Q61L were incubated with MCF10A cell extracts, then subjected to GST pull down. Western blots with the indicated antibodies. For both experiments, 3 biological repeats with similar results, only one is displayed. n corresponds to the total number of beads quantified in all 3 experiments. ****P<0.0001.

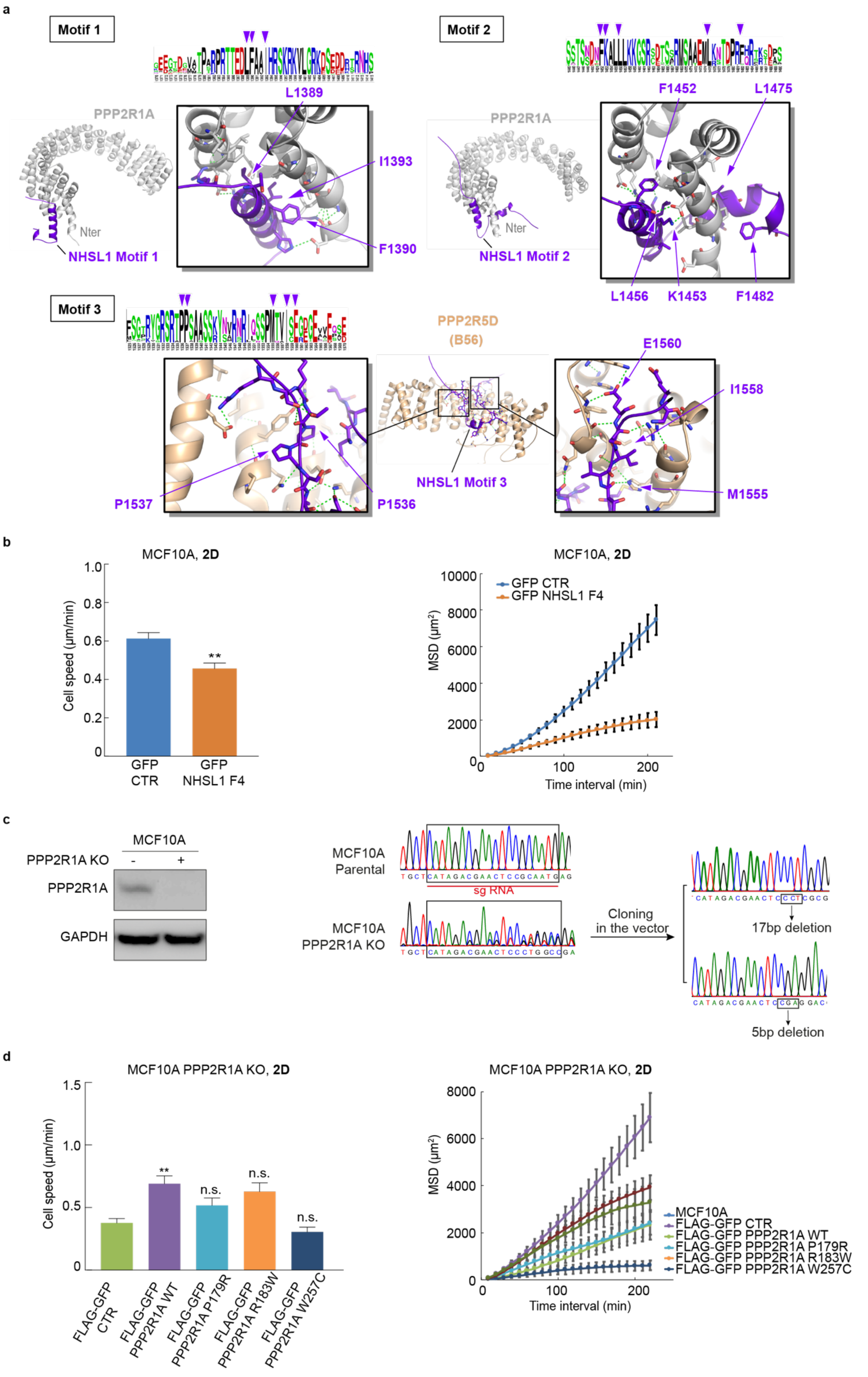

a
Motif 1
L1389
PPP2R1A
I1393
F1390
Nter
NHSL1 Motif 1
Motif 2
F1452
L1475
PPP2R1A
Nter
NHSL1 Motif 2
L1456
K1453
F1482
Motif 3
PPP2R5D
(B56)
E1560
I1558
NHSL1 Motif 3
P1537
P1536
M1555
b
MCF10A, 2D
Cell speed (µm/min)
GFP CTR
GFP NHSL1 F4
**
MCF10A, 2D
MSD (µm²)
GFP CTR
GFP NHSL1 F4
Time interval (min)
c
MCF10A
PPP2R1A KO
PPP2R1A
GAPDH
MCF10A Parental
sg RNA
MCF10A PPP2R1A KO
Cloning in the vector
17bp deletion
5bp deletion
d
MCF10A PPP2R1A KO, 2D
Cell speed (µm/min)
**
n.s.
n.s.
n.s.
FLAG-GFP CTR
FLAG-GFP PPP2R1A WT
FLAG-GFP PPP2R1A P179R
FLAG-GFP PPP2R1A R183W
FLAG-GFP PPP2R1A W257C
MCF10A PPP2R1A KO, 2D
MSD (µm²)
Time interval (min)
MCF10A
FLAG-GFP CTR
FLAG-GFP PPP2R1A WT
FLAG-GFP PPP2R1A P179R
FLAG-GFP PPP2R1A R183W
FLAG-GFP PPP2R1A W257C

**Figure S5, related to figure 6. Structural motifs of NHSL1 fragment 4 and characterization of the PPP2R1A KO clone. (a)** Prediction of three motifs of NHSL1 by AlphaFold2 that interact with PPP2R1A or PPP2R5D subunits of the PP2A complex. Cartoon representation with interacting residues of NHSL1 highlighted as sticks. The conservation pattern represented in a logo plot using WebLogo[15] indicates the highlighted residues by purple triangles. Displayed complexes are for motif 1, PPP2R1A (grey) and the region $P_{1380}$SRP-DD$H_{1410}$ in NHSL1 (violet); for motif 2, PPP2R1A (grey) and the region $A_{1430}$SP-EP$S_{1490}$ in NHSL1 (violet); for motif 3, PPP2R5D (wheat) and the region $S_{1522}$LS-EP$V_{1569}$ in NHSL1 (violet). **(b)** Cell speed and MSD extracted from cell trajectories displayed in Fig.6c. 3 biological repeats of the same experiment with similar results, only one is displayed. **(c)** Characterization of the PPP2R1A KO MCF10A clone. Western blots of PPP2R1A and GAPDH as a loading control. Both alleles contain a deletion that induces a frameshift (17 bp and 5 bp). **(d)** Cell speed and MSD were extracted from random migration of MCF10A parental cells, *PPP2R1A* knockout cells or KO clones expressing wild type or mutants of PPP2R1A, n=19. Related to the corresponding experiment displayed in Fig.7e. 3 biological repeats of the same experiment with similar results, only one is displayed. ** P<0.01

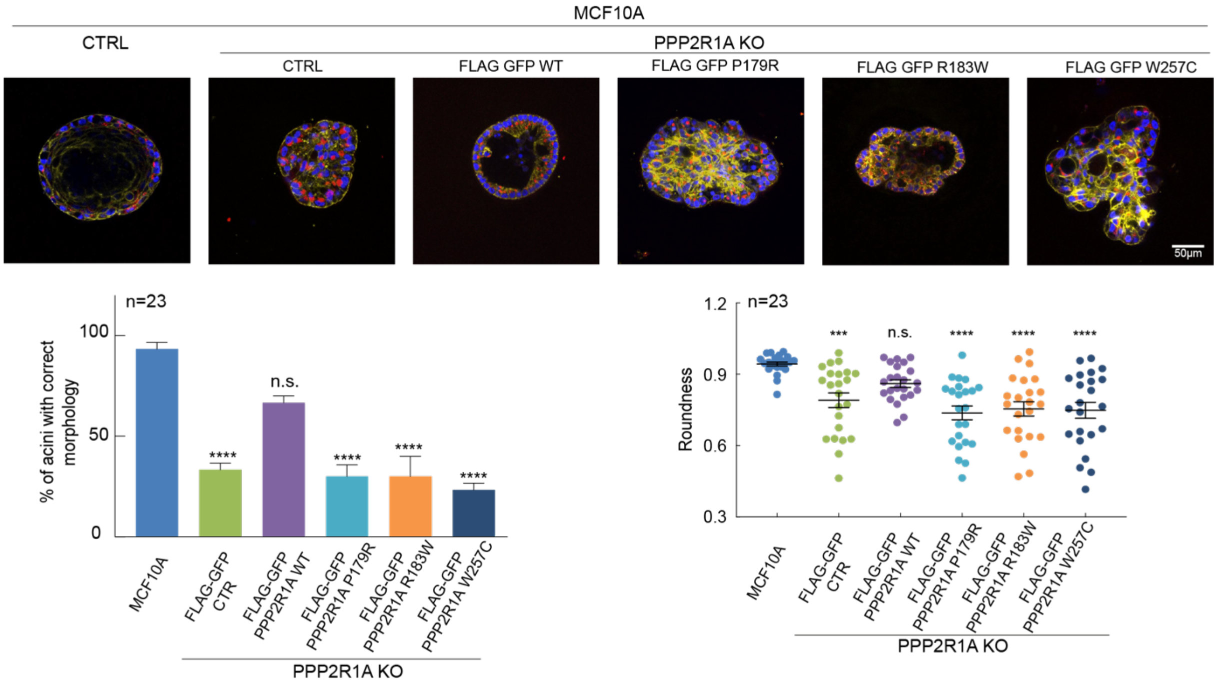


**Figure S6. Development of acini by MCF10A parental cells, *PPP2R1A* KO and rescue derivatives.** Cells were seeded onto Matrigel and grown for 3 weeks. Acini were fixed and stained with DAPI (blue), phalloidin (green) and antibodies targeting the apical Golgi marker GM130 (red). Multicellular structures were observed using confocal microscopy. 3 biological repeats with similar results. Acini with altered polarity or no lumen were scored as abnormal. Roundness of acini outlines, based on central confocal sections. Quantification of the 3 biological repeats. n=23. ***P<0.001; ****P<0.0001

#### MOVIE LEGENDS

**Movie S1, related to figure 1d. Effect of PPP2R1A depletion on the migration persistence of MCF10A cells.**
MCF10A cells transfected with pools of control (CTRL) or PPP2R1A siRNAs were recorded and tracked. Scale bar: 40µm.

**Movie S2, related to figure 1e. Effect of PPP2R1A depletion on the migration persistence of MCF10A RAC1 Q61L cells.**
MCF10A RAC1 Q61L cells transfected with pools of control (CTRL) or PPP2R1A siRNAs were recorded and tracked. Scale bar: 40µm

**Movie S3, related to figure 1f. Effect of PPP2R1A overexpression on the migration persistence of MCF10A cells.**
MCF10A cells stably expressing FLAG-GFP or FLAG-GFP PPP2R1A were recorded and tracked. Scale bar: 40µm.

**Movie S4, related to figure 1g. Effect of PPP2R1A depletion on the migration persistence of MDA-MB-231 cells in 3D.**
MDA-MB-231 cells transfected with pools of control (CTRL) or PPP2R1A siRNAs were seeded in 3D collagen gels and recorded. Scale bar: 40µm.

**Movie S5, related to figure 3a. Localization of PPP2R1A and NHSL1 in B16-F1 cells.**
B16-F1 cells transiently transfected with mScarlet-PPP2R1A and GFP-NHSL1 were recorded. Scale bar: 10µm.

**Movie S6, related to figure 3c. Localization of PPP2R1A and ARPC1B in B16-F1 cells.**
B16-F1 cells transiently transfected with mScarlet-PPP2R1A and GFP-ARPC1B were recorded. Scale bar: 10µm.

**Movie S7, related to figure 4a. Effect of PPP2R1A and NHSL1 combined depletion on the migration persistence of MCF10A cells.**
MCF10A cells transfected with indicated siRNA pools were recorded and tracked. Scale bar: 40µm.

**Movie S8, related to figure 4b. Effect of PPP2R1A and NHSL1 combined depletion on the haptotaxis of MCF10A cells along the fibronectin gradient.**
MCF10A cells transfected with indicated siRNA pools were recorded and tracked. Scale bar: 40µm.

**Movie S9, related to figure 6c. Effect of NHSL1 fragment 4 on the migration persistence of MCF10A cells.**
MCF10A cells stably transfected with GFP or GFP-NHSL1 fragment 4 were recorded and tracked. Scale bar: 40µm.

**Movie S10, related to figure 6f. Effect of PPP2R1A mutations on the migration persistence of MCF10A cells.**

*PPP2R1A* knockout cells were generated in MCF10A. Then the knockout cells were stably transfected with WT or mutant forms of PPP2R1A. Cells for each condition were recorded and tracked. Scale bar: 40µm.
